# Face-selective cortical regions inherit the visuospatial organisation of early visual cortex

**DOI:** 10.1101/2024.05.14.594166

**Authors:** Anisa Y. Morsi, Hugo T. Chow-Wing-Bom, D. Samuel Schwarzkopf, Valérie Goffaux, Tessa M. Dekker, John A. Greenwood

**Author notes:** Corresponding author: John A. Greenwood, Address: UCL Experimental Psychology, 26 Bedford Way, London WC1H 0AP.

## Abstract

Face recognition relies on dedicated brain regions that are widely considered to show unique selectivity, including a disproportionate vulnerability to face inversion and a relative invariance to stimulus location. This proposed spatial invariance contrasts with accounts of common ‘visuospatial coding’ whereby high-level category-selective areas inherit spatial properties from earlier regions. Critically, early regions (V1-V3) show characteristic retinotopic variations, with greater cortical sampling along the horizontal than vertical meridian and in the lower than upper field, mirroring established behavioural advantages for face recognition at these locations. We examined whether face-selective regions (OFA, pFus, mFus) share these spatial anisotropies, and whether these properties could drive the observed variations in face recognition. Using wide-field retinotopic mapping with upright and inverted faces (±21° eccentricity), we estimated population receptive fields (pRFs) and visual-field coverage. Though pRFs were substantially larger in face-selective regions than in V1-V3, pRF sizes did not vary in line with behavioural anisotropies. In contrast, both early and face-selective regions showed higher pRF numbers and greater visual-field coverage along the horizontal meridian and in the lower field. These sampling differences provide a plausible neural substrate for behavioural anisotropies in face recognition. We also show that pRF numbers in mFus were greater for upright than inverted faces, likely contributing to the perceptual advantage for upright faces. Together, our findings indicate that variations in visual-field sampling within face-selective cortex parallel those of early visual areas, supporting a hierarchical model in which the spatial selectivity of category-selective areas is built on that of earlier regions.

**Significance statement:** High-level face-selective cortex is often treated as functionally and spatially distinct from early visual areas. We show instead that these regions all share systematic patterns of spatial biases. Using population receptive field (pRF) analyses, both early and face-selective regions showed greater sampling (higher pRF numbers) and increased visual-field coverage along the horizontal than vertical meridian, and in the lower than upper visual field, matching the pattern of anisotropies in face-recognition performance. This relationship was absent for pRF sizes, indicating that behavioural anisotropies are more closely linked to sampling density and coverage. This common pattern of spatial sampling embeds specialised face-processing systems within a hierarchical network in which high-level regions retain fundamental aspects of visuospatial organisation from early visual cortex.

## Introduction

Face recognition is essential to everyday life, though its complexity places substantial computational demands on the visual system (Bruce and Young, 1986). These pressures have been argued to drive the ‘special’ nature of face processing, supported by a network of specialised brain regions (Kanwisher et al., 1997; Haxby et al., 2000; Grill-Spector et al., 2017). These regions show selectivity for face inversion (Kanwisher et al., 1998) and either complete invariance to the visual-field location of faces (Tanaka, 1996) or distinct patterns of spatial selectivity to early visual cortex (Avidan and Behrmann, 2021; Poltoratski et al., 2021). This apparent uniqueness contrasts with proposals for ‘canonical computations’ in the brain (Miller, 2016) and growing evidence that high-level category-selective regions share a common ‘visuospatial coding’ with earlier areas, matching the statistical properties of the visual environment (Groen et al., 2022). We tested this apparent discrepancy by examining whether face-selective regions retain the characteristic retinotopic anisotropies of early visual cortex, with greater resources dedicated to more common object locations, and whether this selectivity explains variations in face perception across the visual field.

A ubiquitous feature of our visual abilities is their variation around the visual field: acuity, contrast sensitivity, and crowding all show better performance on the horizontal than the vertical meridian (horizontal-vertical anisotropy) and in the lower than upper visual field (upper-lower anisotropy; Carrasco et al., 2001; Westheimer, 2003; Abrams et al., 2012; Greenwood et al., 2017; Himmelberg et al., 2020; Barbot et al., 2021). This pattern aligns with the typical locations of behaviourally relevant information in the visual field (Himmelberg et al., 2023a). These performance anisotropies have been linked with variations in retinotopic organisation in early visual cortex. In areas V1-V3, population receptive fields (pRFs) tend to be smaller along the horizontal than vertical meridian (Silson et al., 2018; Silva et al., 2018), though evidence for upper-lower differences in pRF size is less consistent (Silva et al., 2018; Himmelberg et al., 2023b). More consistently, V1 shows greater surface area along the horizontal than vertical meridian and in the lower than upper field (Amano et al., 2009; Himmelberg et al., 2023b). Differential cortical sampling of the visual field could therefore drive the characteristic variations of low-level vision.

Recent behavioural research shows similar anisotropies for face perception, with better recognition along the horizontal than vertical meridian and in the lower than upper field (Roux-Sibilon et al., 2023; Morsi et al., 2024). Whether these anisotropies arise from face-selective brain areas is unclear. Despite early accounts that face-selective areas are invariant to visual-field location (Tanaka, 1996), recent studies reveal clear retinotopic organisation: pRF sizes increase with eccentricity, with over-representation of the fovea (Kay et al., 2015; Finzi et al., 2021; Silson et al., 2022), as in earlier areas (Amano et al., 2009). However, the ‘uniqueness’ of higher-level face-selective regions has been proposed to extend to this retinotopic selectivity. Namely, the superior recognition of upright faces (Yin, 1969; Rossion and Gauthier, 2002) has been linked with *larger* pRFs compared with inverted faces (Witthoft et al., 2016; Poltoratski et al., 2021), leading to proposals that enhanced upright-face recognition is driven by larger pRFs and associated increases in visual-field coverage (Avidan and Behrmann, 2021; Poltoratski et al., 2021). Such a relationship contrasts with early visual cortex, where smaller receptive fields are associated with better acuity (Duncan and Boynton, 2003; Silva et al., 2021).

These findings highlight a puzzling dissociation between low– and high-level vision. One account posits that face-selective regions inherent canonical tuning properties from early cortex, with greater spatial precision and/or increased neural sampling along the horizontal than vertical meridian and the lower than upper field in all regions. The other proposes that the specialisation of face-selective regions gives a unique spatial selectivity, with either invariant or reversed tuning properties relative to earlier areas. To address this, we used high spatial precision pRF mapping with upright and inverted face stimuli to characterise spatial tuning across the visual hierarchy.

## Methods

### Design

To compare the spatial selectivity of early– and face-selective cortical regions, we undertook retinotopic mapping procedures with wide-field bars containing either upright or inverted face stimuli. Visual-field variations were measured for three retinotopic properties: pRF size, to quantify the spatial tuning aperture; pRF numbers, to quantify the size of the neural population sampling a given retinotopic location; and visual-field coverage, which combines these properties to quantify the total sampling resource dedicated to the visual-field location. In this way, we compared early visual areas V1-V3 with core regions of the face-processing network: the occipital face area (OFA), and the posterior (pFus) and medial (mFus) fusiform gyrus which comprise the Fusiform Face Area (FFA). If face-selective regions share their spatial selectivity with earlier regions V1-V3 (Groen et al., 2022), these properties should vary similarly, with e.g. smaller pRFs, higher pRF numbers, and greater coverage along the horizontal vs. vertical meridian and in the lower vs. upper field. Alternatively, if the spatial selectivity of face-selective regions reflects specialised mechanisms, whereby large receptive fields benefit face perception (Witthoft et al., 2016; Avidan and Behrmann, 2021; Poltoratski et al., 2021), we might instead observe either wholly unique variations or a reversal of the spatial selectivity in face-selective regions. To further test the link between this spatial selectivity and the stimulus selectivity of face-selective regions (i.e. the face inversion effect), we also measured these retinotopic properties with both upright and inverted face stimuli, seeking to replicate prior findings that pRF sizes are larger and visual-field coverage greater for upright vs. inverted faces (Poltoratski et al., 2021). Together, this design allowed us to assess whether the retinotopic properties of face-selective regions vary across the visual field and with stimulus properties in a manner that could drive observed variations in face perception.

### Participants

Ten participants (six female, M_age_ = 29.1 years) took part, all of whom had normal or corrected-to-normal vision and gave written informed consent. Procedures were approved by the Experimental Psychology Research Ethics Committee at University College London.

### Apparatus

Functional and anatomical scans were obtained using a Prisma 3T MRI scanner (Siemens, Erlangen, Germany). Stimuli were displayed on a back-projection screen in the bore of the magnet using an EPSON EB-L1100U projector that had a maximum luminance of 502 cd/m^2^. The projector had a refresh rate of 60 Hz and a resolution of 1920 x 1200 pixels, with stimuli displayed in the central 1200 pixels (at a physical size of 27 x 27 cm). Participants viewed the screen at a distance of 34 cm via a mirror mounted above the head (see Chow-Wing-Bom et al., 2025), giving a maximum field of view of 43.3° (±21.65° eccentricity). Gamma correction was performed, with the grey stimulus background presented at the mean projector luminance (251 cd/m^2^).

### Stimuli

Where prior work has examined the spatial selectivity of face-selective cortex by presenting one face at a time (Poltoratski et al., 2021), we developed totem-pole style bars of faces to more closely match the stimuli used for retinotopic mapping more broadly. While there is evidence that wedge-and-ring stimuli give pRF estimates with better goodness-of-fit in V1 compared to bars (Alvarez et al., 2015), the use of wedge-and-ring stimuli requires eccentricity scaling to an extent that is unclear in face-selective regions. Instead we used bars, which have been shown to yield more accurate estimates of pRF eccentricity than wedge-and ring-stimuli (Linhardt et al., 2021). Where prior studies have examined only small regions of the visual field, here we wanted to cover a wider expanse of the visual field, particularly given the large sizes of pRFs in face-selective regions. Our bars thus covered a large field of view in length (43.3°; Figure 1A), with both horizontal and vertical orientations.

**Figure 1.**
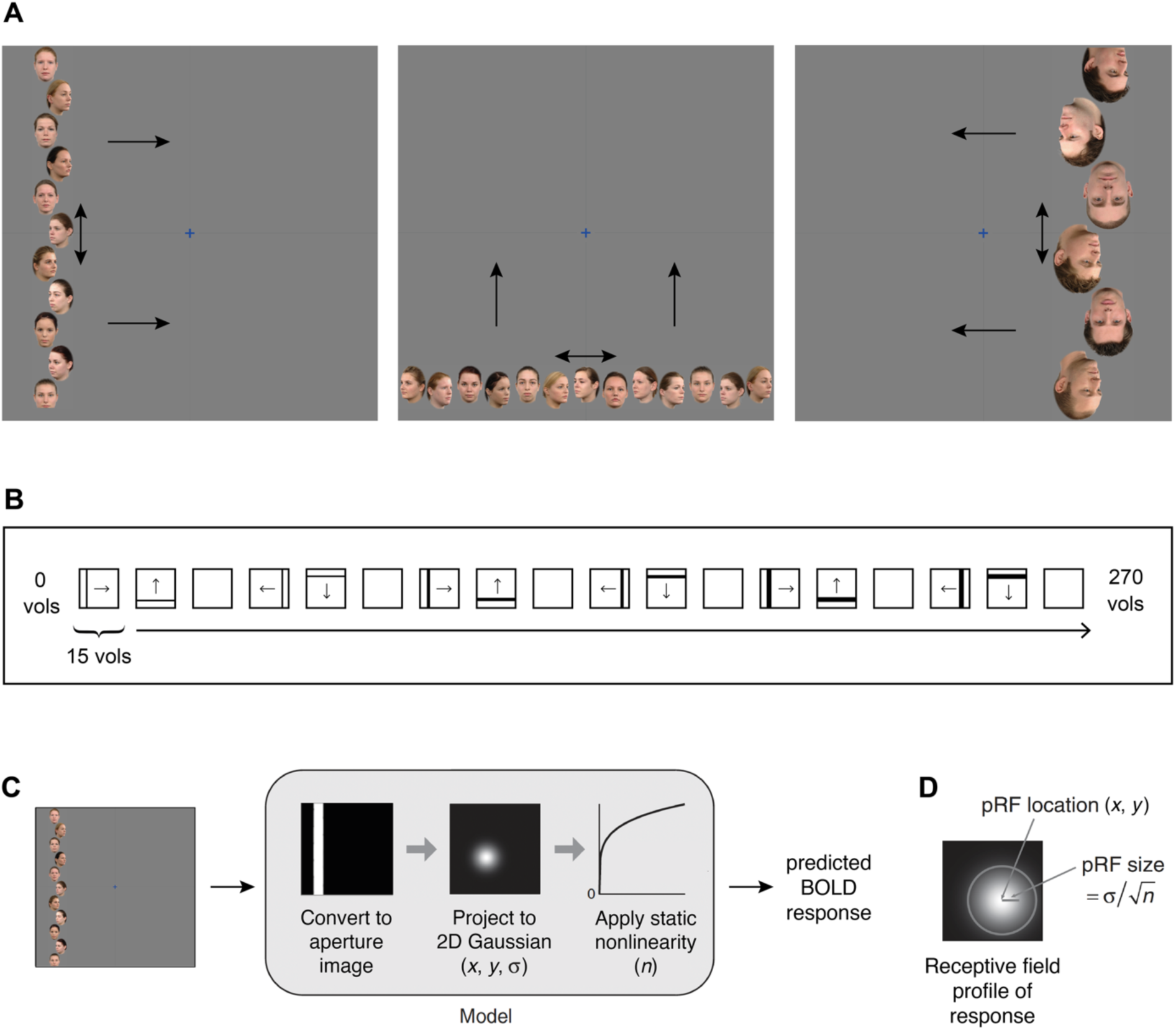
A. Example retinotopic mapping stimuli. A blue fixation cross appeared at the display centre, while bars containing either male or female faces traversed the screen in one of four cardinal directions (shown by unidirectional arrows). Bars appeared at one location per TR (one second), moving back-and-forth along the length of the bar (shown by bidirectional arrows). Different bar widths and face orientations are shown in each panel. **B.** Bar conditions throughout the experiment. Each square represents one sweep across the screen (with 15 locations per sweep). The narrowest bars were shown first, proceeding to the widest. Blank periods (15 TRs) occurred after every two sweeps. Arrows represent the bars’ direction of movement. **C.** pRF model. The stimulus was converted to a binary aperture image, with each pRF modelled as a 2D Gaussian before a static nonlinearity was applied using a compressive spatial summation parameter. The model output gives the predicted BOLD response. **D.** A depiction of pRF location and size in the compressive spatial summation model. Position is determined by *x* and *y* coordinates, while size is the standard deviation (*α*) divided by the square root of the spatial summation exponent *n*, adapted from Kay et al., (2013).

Stimuli were programmed using MATLAB (MathWorks, Inc) and PsychToolbox (Brainard, 1997; Pelli, 1997; Kleiner et al., 2007). Faces in each bar were selected from 15 male and 15 female faces with a neutral expression, taken from the Radboud Face Database (Langner et al., 2010) and presented in colour. To maximise face-selective activation, we sought to minimise crowding between the faces (Louie et al., 2007; Kalpadakis-Smith et al., 2018), adaptation (Fang et al., 2007), and repetition suppression effects (Louie et al., 2007; Henson, 2016) by using faces with three viewpoints according to the view of the model: front-(90°), left– (135°) and right-facing (45°). This resulted in a total of 90 face images, which had their background removed and were resized to 332×450 pixels using Adobe Photoshop CS6. To further avoid adaptation/suppression effects tied to face identity (Natu et al., 2016), each identity could only appear once in a given bar. For the inverted face bars, faces were flipped along the vertical axis (Kalpadakis-Smith et al., 2018). The background of each bar matched the grey background of the experimental screen.

To ensure the visibility of faces in the periphery, and to improve pRF fitting by maximally activating differently sized pRFs (both smaller pRFs near the fovea and larger pRFs peripherally), three bar widths were used: 5.3°, 7.0° and 10.1°. Face size was determined by the bar width (i.e. faces scaled with bar sizes). For each bar width, bar orientation (horizontal and vertical) and face orientation (upright or inverted), five bars containing male faces and ten bars containing female faces were generated (male bars appeared less frequently, as described below). Each bar contained faces of different viewpoints in a pseudo-randomised manner, such that faces of one viewpoint could not appear next to a face of the same viewpoint.

To fill each bar, faces were shifted along the width of the bar (the x-axis for vertical bars and the y-axis for horizontal bars) so that they could be moved closer together along the opposite axis, reducing blank space. To further maximise the activation of face-selective regions and ensure that the time-averaged bar locations contained faces in as much of the bar as possible, faces were moved along the length of the bar during presentation. To produce this motion, bars were made longer than required and cropped to screen dimensions such that faces at the extreme edges moved off-screen at times. Displaced versions of each bar image were shown for 66.7 ms each to give motion at a speed of 13.65, 17.63, or 25.59°/s (increasing with the width of the bars/faces). As such, faces within the bars moved smoothly side to side (horizontal bars) or up and down (vertical bars) at each location.

### Procedure

Each run of the retinotopic-mapping procedure began with a blank screen for five seconds, with a central fixation cross subtending 0.95° of visual angle. Bars then stepped across the screen in four directions: 0° (rightwards), 90° (upwards), 180° (leftwards) and 270° (downwards), with one location per repetition time (TR), which lasted one second (Figure 1B). As described above, faces moved along the length of the bar in each location to ensure that the time-averaged bar for each TR contained faces across the whole bar. Each sweep across the screen contained 15 equal steps, meaning that steps were smallest with the largest bar widths and vice versa. The number of steps was matched to avoid the pRF fitting being biased towards bar widths with more TRs (by contributing more to the least-squared error between data and model predictions used for model fitting). The thinnest bars were presented first (four sweeps, one per direction of motion) before moving on to the next thickness (Figure 1B). As there were three bar thicknesses, each run had a total of 12 sweeps. Every second sweep was followed by a blank period of 15 TRs. Each run therefore comprised 270 TRs.

Participants were instructed to maintain fixation on a cross at the screen centre, whilst performing two tasks. To ensure fixation, participants reported when the fixation cross changed from blue to purple (0.002 probability, and lasting 0.2 seconds). To ensure that attention was directed towards the bars at the same time, participants also responded when a bar containing male faces appeared in a given TR. Most bars contained female faces, while bars consisting of male faces occurred with 0.075 probability. Responses were recorded via a button box. Participants did not receive feedback, however key presses were monitored throughout the experiment to ensure that participants were engaging with the task. Upright and inverted runs were interleaved to avoid effects like fatigue disproportionately affecting one condition.

### Localisation of face-selective regions

To identify face-selective Regions of Interest (ROIs), a functional localiser was run in the same scan session. In prior studies, these localisers have tended to use full-field stimuli presented foveally, with the subsequent investigation of spatial selectivity in face-selective regions then using stimuli presented only in the central 6-8° of the visual field. In scaling stimuli up to our 43.3° field of view, we were concerned that this approach would sub-optimally drive face-selective regions, given that large faces are less effective at engaging holistic processes (McKone, 2009). A second concern was that the use of full-field stimuli shown foveally may bias the localisation of face-selective regions towards voxels with a preference for large foveal stimuli, thereby exaggerating the extent of the foveal bias within face-selective regions.

For these reasons, we developed a novel localiser which presented faces and objects in both foveal and peripheral locations at a range of sizes. Images of faces, hands and instruments were displayed to cover the full 43.3° field of view. To maximise both foveal and peripheral stimulation, two configurations of stimuli were used: large, single images centred on the fovea, and smaller images tiled across the screen in a 3×3 grid (Figure S1). Similar to existing localisers (Stigliani et al., 2015; Schuurmans et al., 2023), faces, hands, and instruments were used as stimuli to isolate face-selective regions. Twenty images were created for each category, with contrast normalisation applied to give images a root mean square (RMS) contrast of 0.15. Twenty tiled images for each object category were then created by randomly selecting 9 images from the same category for the grid, ensuring that the same image did not appear twice. This gave 3 object categories (faces, hands, instruments) and 2 tiling conditions (single or tiled). In each case, images were shown on a noise background produced by iterative phase scrambling – each image (or image set) underwent a fast Fourier transform (FFT) followed by phase scrambling, pasting the faces/objects back onto this scrambled image, and scrambling again, with 500 repetitions (Petras et al., 2019).

Images from each of the 6 conditions were shown in blocks lasting 10 seconds each, with 20 stimuli from a given condition displayed consecutively for 500 ms each with no inter-stimulus interval. Single and tiled configurations were presented in separate blocks. Each run consisted of 51 blocks, with baseline (blank) periods for 10 seconds to begin and end. Participants were instructed to maintain fixation and press a button when a phase-scrambled image appeared. Each participant completed two runs.

To identify face-selective brain regions, we contrasted blood-oxygen-level-dependent (BOLD) responses to faces against the other object categories (Kanwisher et al., 1997; Weiner and Grill-Spector, 2010), combining both single and tiled versions (single and tiled faces vs. single and tiled hands, plus single and tiled instruments) using a general linear model (GLM) performed in SPM12 (Penny et al., 2011). Statistical contrasts were carried out using a threshold of *t* ≥ 2, chosen to maximise the number of pRFs remaining for further analyses after filtering by visual field location. We defined three face-selective areas (OFA, pFus and mFus; see Figure 2A for an example) in nine participants, though pFus could not be defined in one participant. Statistical *T* maps were surface projected using Freesurfer (Fischl, 2012) and used as a visual guide during the delineation of face-selective ROIs. Large areas were initially drawn manually, before an automatic process was used to define the ROI by identifying the vertex with the peak *T* statistic in each region and incorporating neighbouring vertices that were above the *T* threshold (*t* ≥ 2). The inclusion of tiled images gave a significant increase in the vertices within each ROI compared to the use of single images alone (Figure S2).

**Figure 2.**
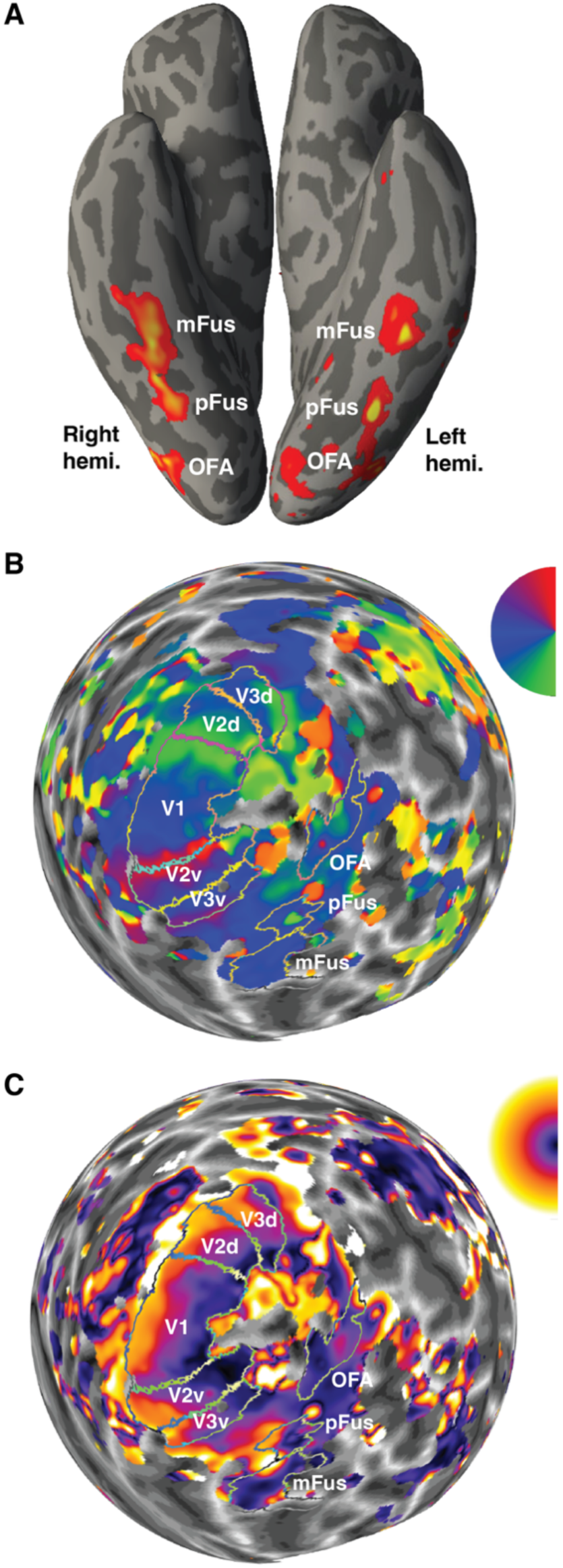
A. The location of face-selective ROIs (OFA, pFus, and mFus) for an example participant. Activation maps (determined by statistical contrasts on localiser runs) are shown on the ventral view of an inflated 3D model (with left and right hemispheres indicated). Smoothing has been applied for visualisation, which was not used for the statistical contrasts. **B.** An example retinotopic map of polar angle preferences from the right hemisphere of one participant, plotted on an inflated, spherical cortical surface. Delineations of V1-V3 and OFA, pFus and mFus are outlined. The colour wheel indicates polar angle coordinates (green for the lower visual field, blue around the horizontal meridian, red for the upper field). **C.** An eccentricity map for the same participant, plotted similarly to panel B, where purple represents central eccentricities and yellow more peripheral locations up to 21.65°.

### MRI data acquisition

A 64-channel head coil was used with the 3T scanner, with cushions to minimise head movement. Functional scans were run with only the back of the head coil to increase the field of view, leaving 42 channels. A T1-weighted anatomical magnetisation-prepared rapid acquisition with gradient echo (MPRAGE) image was acquired (TR = 2300 ms and TE = 2.98 ms, voxel size = 1 mm isotropic voxels), along with functional T2-weighted multiband 2D echoplanar images (TR = 1000 ms, TE = 35.20 ms, voxel size = 2 mm isotropic voxels, 48 slices, flip angle = 60°, acceleration factor = 4). Each functional scan contained 270 volumes. A 30 second localiser was carried out before the functional scans and again before the anatomical scan, after the front head coil was fitted. Fixation was monitored throughout the experiment using an Eyelink 1000, though we did not record fixation data.

### MRI data preprocessing

For each participant, the T1 anatomical scan was automatically segmented, with the boundary between grey and white matter used to generate an inflated 3D representation of the cortical surface using Freesurfer 7.1.1 (Dale et al., 1999; Fischl et al., 1999; Fischl, 2012). Functional images were B0 distortion corrected and motion corrected using AFNI software (Cox, 1996). An alignment volume was created by finding the volume with the fewest voxel outliers across all runs, which all functional volumes were then aligned to. Using Freesurfer (Fischl, 2012), the alignment volume was co-registered to the structural image, and surface projection was performed.

### pRF fitting

The four retinotopic-mapping runs were averaged and pRF analyses carried out using the SamSrf 9.4 MATLAB toolbox (Schwarzkopf, 2022). Following recent examinations of face-selective pRFs (Poltoratski et al., 2021), and demonstrations that stimulus responses in category-selective regions sum in a sub-additive rather than linear manner (Kay et al., 2013), a Compressive Spatial Summation (CSS) model (Kay et al., 2013) was used to estimate pRF parameters. This was implemented within SamSrf 9.4 (Figure 1C), with each pRF estimated as a two-dimensional Gaussian before a compressive non-linearity was applied. The CSS model fitting involved four free parameters: *x* and *y* (the position of the pRF centre within the visual field), σ (the standard deviation or spatial spread of the pRF, in degrees of visual angle) and *n* (the exponent of the compressive non-linearity; Figure 1D). As the compressive nonlinearity affects the spread of the receptive field profile, during analyses we defined pRF size as σ divided by the square root of the exponent *n* (Kay et al., 2013). Stimulus locations were fed into model fitting via stimulus apertures created for each run, which were averaged across the four runs, resulting in one set of apertures comprising 270 frames (one for each TR). Because the faces moved within the bar stimuli (as described above), averaging these stimulus positions over time meant that the apertures formed solid bars, similar to standard retinotopic mapping procedures (Figure 1C).

A coarse-to-fine approach was used for pRF fitting. For each vertex, the coarse fit was carried out using an extensive multidimensional search space of 35,496 grid points, with different combinations of *x*, *y* and σ. The parameters with the highest Pearson correlation between the predicted and observed time series were then selected. These values were used to seed the fine fit, which used the Nelder-Mead simplex-based method (Nelder and Mead, 1965) to reduce the residual sum of squares (RSS) between the predicted and observed time series, and determine optimal values for the four free parameters (*x*, *y*, σ, and *n*).

### Delineation of ROIs

Prior to delineation, vertices with a goodness-of-fit threshold below 0.2 were removed, and a smoothing kernel of 3 mm full width half maximum (FWHM) was applied. pRF locations (*x* and *y*) were then used to project colour-coded polar angle and eccentricity maps onto the cortical surface (see examples in Figure 2B-C). Early visual areas V1-V3 were delineated using an auto-delineation tool and then corrected manually using SamSrf 9.4 (Schwarzkopf, 2022). This involved using standard criteria based on reversals in polar angle (DeYoe et al., 1994; Sereno et al., 1995; Engel et al., 1997), assisted by the eccentricity maps. These early regions were delineated using the maps for the upright face condition, before being checked and adjusted (if needed) using the inverted maps to ensure correspondence between the two. Maps in early visual areas were nonetheless highly consistent between runs with upright and inverted faces. Face-selective areas were delineated via localiser analyses, as above.

### Vertex selection

To avoid artefacts, vertices that had beta amplitudes below 0.01 or above 3 (*z* scores), sigma values of 0, or which were located perfectly at the centre (*x* and *y* of exactly 0, indicative of fitting errors) were removed. To avoid noisy and unreliable vertices, those with a goodness-of-fit threshold (*R^2^*) below 0.2 were also removed (as above). In OFA and mFus, some participants showed vertices with very small pRF size estimates (almost 0) at high eccentricities, which upon closer analysis were the result of poor fits. To avoid these unreliable estimates affecting the main pattern of results, we increased the *R^2^* threshold within face-selective ROIs for some participants (OFA: four participants = 0.4, mFus: one participant = 0.4, one participant = 0.3).

### Location analyses

To compare pRF properties across the visual field, pRFs were filtered according to their centre position. Four quadrant (90°) wedges were defined, each of which included polar angle locations within ±45° on either side of the left horizontal, right horizontal, upper vertical and lower vertical meridians. Although behavioural research has suggested that visual field anisotropies decline at locations more than 30° away from the meridian (Abrams et al., 2012; Benson et al., 2021), fMRI studies have found similar asymmetries in cortical surface area across different wedge widths (Himmelberg et al., 2023b), and that anisotropies in pRF properties were evident with 90° wedges (Silva et al., 2018). In our data, patterns were similar with wedge widths of 60° and 90°. We thus used the wider 90° width, which retained a higher number of pRFs after filtering. For horizontal-vertical comparisons, the left and right wedges were combined to make the horizontal meridian, with upper and lower wedges combined to make the vertical meridian.

### Visual-field coverage

Visual-field coverage was calculated by generating a Gaussian receptive field profile for each vertex based on its centre position (*x,y*), eccentricity, and sigma (σ), and then raising the receptive field profile by the spatial summation exponent (*n*), matching the best-fitting CSS model. Receptive field profiles were then summed across vertices to give coverage values across the visual field for each ROI, face orientation (upright/inverted), and participant. Because absolute values would differ based on numerous factors (e.g. the number of vertices, sigma values, and exponents), we normalised coverage values by dividing by their maximum value across both upright and inverted maps (separately for each ROI and participant). Coverage therefore represents the proportion of the peak response (within each ROI) at each visual field location. Coverage values were extracted from these plots according to eccentricity and polar angle location (using the wedges described above) for further analyses. Note that some prior studies have calculated coverage using binary circles (e.g. Witthoft et al., 2016; Poltoratski et al., 2021). Although these binary approaches incorporate the position and size of receptive fields, they do not capture the spatial profiles of the constituent pRFs. By summing the profiles, we aimed to better account for the spatial pattern of responsiveness within each pRF when generating estimates of coverage (as in e.g. Amano et al., 2009).

### Statistical analyses

We were interested in three pRF properties: size 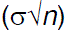, number (the total amount of pRFs after poor-fitting vertices were removed) and visual-field coverage (the values extracted from the coverage plots). These properties were examined in eccentricity bins of 1° width, ranging from from 0.5° to 21.5°.

Linear mixed effects models were used to investigate whether location, inversion and eccentricity could predict pRF size. Because the location of each pRF was determined by its centre, size could not be estimated if there were no pRF centres within that region. Linear mixed models can deal with these ‘missing’ estimates by examining the linear change in pRF size with eccentricity. Separate mixed effects models were run for each ROI (V1, V2, V3, OFA, pFus, and mFus). Our main analyses examined pRF size with fixed factors for visual field location (horizontal/vertical or upper/lower), eccentricity and inversion (upright/inverted), with a second analysis examining fixed factors for eccentricity and inversion (upright/inverted) irrespective of location. Participant was specified as a random factor for the intercept as well as for each of the fixed factors, as the slope of the relationship between pRF size and eccentricity, location and/or inversion could vary across individuals. Differences at each eccentricity were then examined using Wilcoxon signed rank tests.

Because pRF number and visual-field coverage showed non-linear profiles, mixed effects analyses of variance (ANOVAs) were used to assess the effects of eccentricity, inversion, location and participant on these properties. Separate ANOVAs were run for each ROI (V1, V2, V3, OFA, pFus and mFus). Our main analyses included within-subjects fixed factors for location, eccentricity, and inversion (upright/inverted), with a second analysis run to examine effects of inversion regardless of location. In all ANOVAs, participant was entered as a between-subjects random factor. Inversion effects and location differences were then examined via t-tests or Wilcoxon signed rank tests (if sphericity or homoscedasticity assumptions were violated).

## Results

We used population receptive field (pRF) mapping to compare the spatial selectivity of early visual cortex (V1-V3) with three face-selective regions of ventral occipitotemporal cortex (OFA, pFus, mFus). Best-fitting parameters for the CSS model gave a good characterisation of the BOLD responses to our stimuli, with high average R^2^ values and equivalent levels with both upright and inverted face stimuli (Figure S2A). Consistent with prior work (Kay et al., 2013; Poltoratski et al., 2021), the exponent of the compressive non-linearity (the *n* parameter) averaged below 1 in all regions tested (indicating a compressive nonlinearity), and decreased in the higher face-selective regions of the visual hierarchy (Figure S2B). Maps of the full visual-field coverage for each ROI are displayed in Figure 3, separately for upright and inverted faces. All ROIs showed some degree of coverage throughout the visual field, including at the farthest eccentricities tested, though a bias towards foveal locations is also evident, particularly in face-selective regions. While coverage patterns are similar regardless of face orientation in early visual cortex, in mFus there is broader coverage in both peripheral and central locations for upright vs. inverted faces.

**Figure 3.**
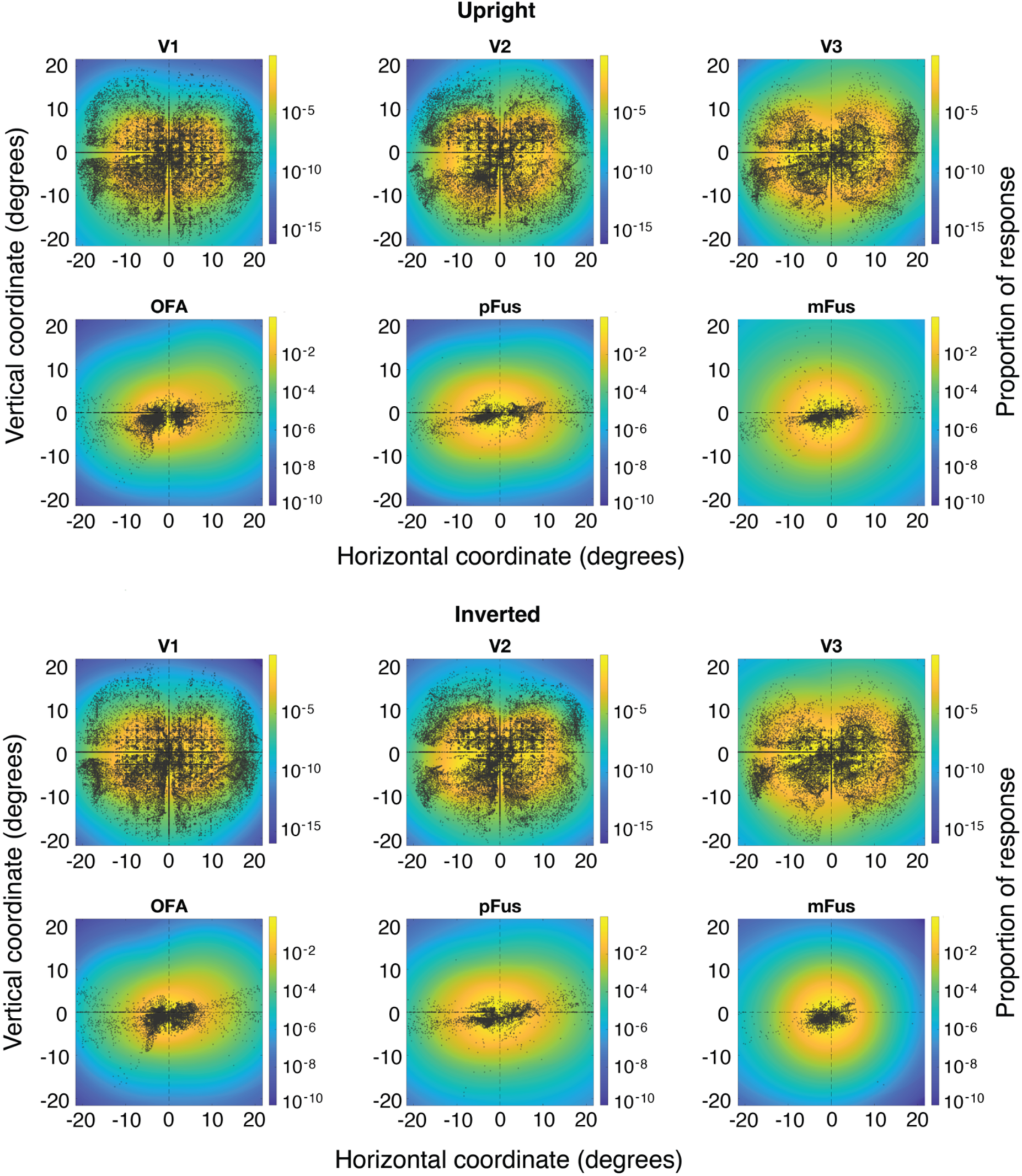
Mean visual-field coverage for upright (top) and inverted (bottom) faces, plotted separately for early (V1-V3) and face-selective (OFA, pFus, mFus) cortical regions. Coordinates represent visual field position in degrees of visual angle. Coverage values were converted to log units for visualisation purposes (see colour bar). Dots represent pRF centres pooled across all participants. These plots show the relative responsiveness of pRFs across the visual field, incorporating the number, size, and spatial profile of each pRF within the population of vertices that responded to each stimulus.

We first assessed how retinotopic properties (pRF size, number, and visual-field coverage) differed according to polar angle, and in particular whether variations in this *spatial selectivity* follow the expected anisotropies along the horizontal vs. vertical meridian and in the upper vs. lower field. As we observed similar location-based variations in these measures for upright and inverted faces, the following sections on visual field anisotropies discuss the results for upright faces only. We then examined whether variations in these retinotopic properties might also subserve the *stimulus selectivity* of face perception for upright rather than inverted faces.

### Visual field anisotropies

#### pRF size

Consistent with prior work, pRF sizes increased throughout the visual hierarchy. Sizes also increased with eccentricity in all visual regions, as shown in Figure 4. We first compared the horizontal and vertical meridians (Figure 4A). In early visual cortex, pRF sizes did not differ significantly along the horizontal and vertical meridians in V1 (*β* = 0.13 [-0.16, 0.44], p = .373) nor V2 (*β* = –0.01 [-0.38, 0.36], *p* = .953), contrary to predictions. A main effect of location was found in V3 (*β* = 0.79 [0.17, 1.41], *p* = .012), with Wilcoxon tests indicating that pRFs were significantly larger along the vertical than horizontal meridian at eccentricities from around 5-15° (see lines in Figure 4A), consistent with the pattern predicted by behavioural anisotropies. In face-selective regions, the number of pRFs in the periphery dropped markedly along the vertical meridian (to be discussed below regarding pRF number), meaning that size estimates were missing at many eccentricities. However, pRF sizes did not differ significantly between the horizontal and vertical meridian in any of these areas, neither for the OFA (*β* = –0.48 [-2.05, 1.10], *p* = .551), pFus (*β* = 0.24 [-1.15, 1.63], *p* = .734) nor mFus (*β* = 1.22 [-0.10, 2.54], *p* = .070).

**Figure 4.**
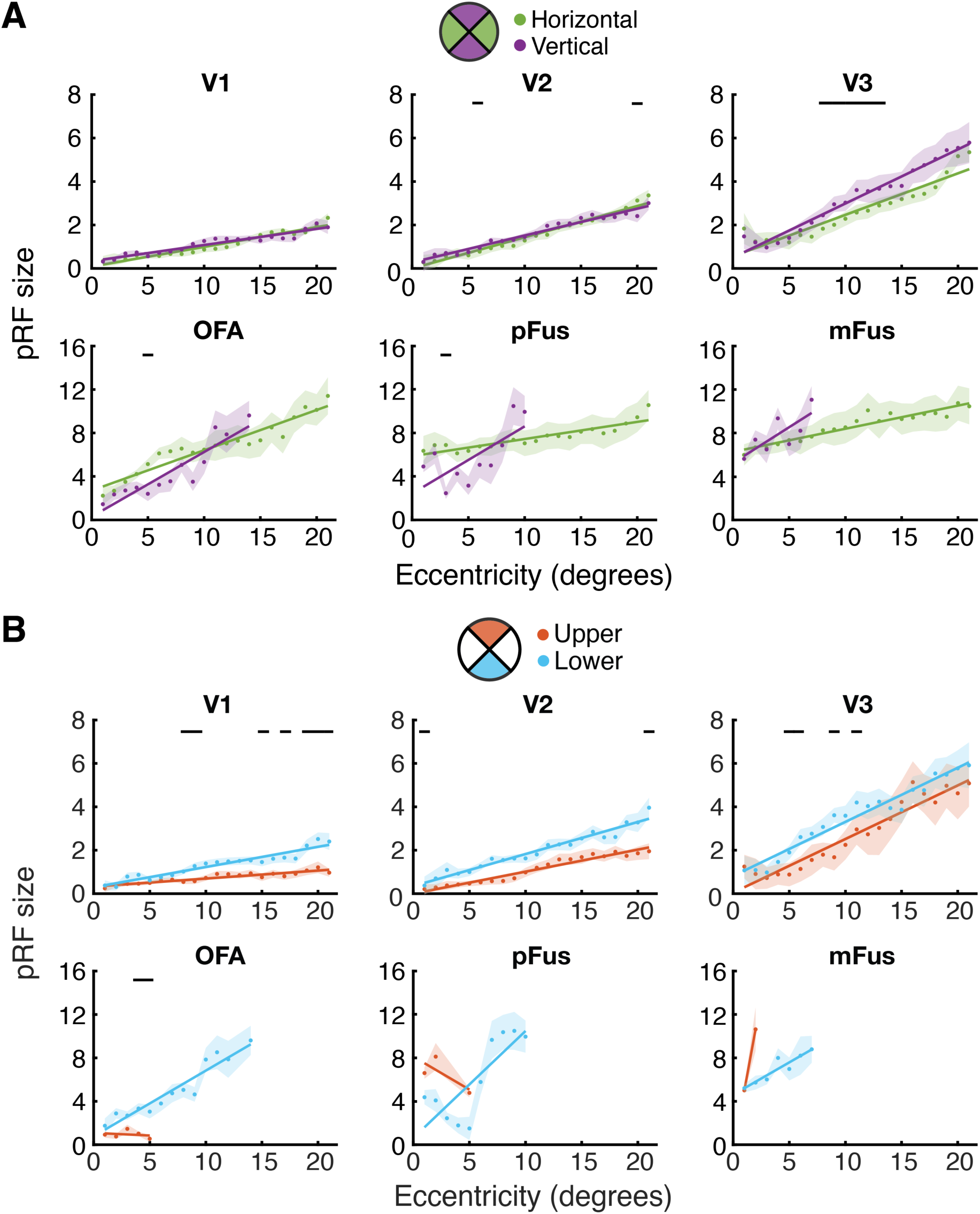
Mean pRF size measured with upright faces across eccentricity along the horizontal (green) and vertical (purple) meridians **(A)** and in the upper and lower visual field **(B)**, plotted separately for early (V1-V3) and face-selective (OFA, pFus, mFus) cortical regions. At each eccentricity, size estimates were only plotted if they were averaged from at least five vertices. Black lines indicate significant differences at each location (*p* < .05). Note differences in the y-axis scale between early and face-selective regions.

Figure 4B plots the comparison of pRF sizes in the upper and lower fields. In early visual cortex, pRFs were larger in the lower than the upper field, with significant main effects of location (V1: *β* = 0.66 [0.20, 1.13], *p* = .006; V2: *β* = 0.88 [0.16, 1.61], *p* = .017; V3: *β* = 0.84 [0.06, 1.62], *p* = .035) and significant t-tests across a range of eccentricities. These differences run in the opposite direction to that predicted by behavioural anisotropies, given that performance is typically worst in the upper field. In face-selective regions, there was also an effect of location in OFA (*β* = 2.30 [0.94, 3.67], *p* = .001), with pRFs again larger in the lower vs. upper field. Size estimates did not differ significantly between the upper and lower field in pFus (*β* = –0.72 [-4.27, 2.83], *p* = .689) or mFus (*β* = –1.72 [-3.76, 0.31], *p* = .096). Taken together, these findings suggest that pRF size was not consistently modulated by location in a manner consistent with behavioural anisotropies, either in early visual cortex or face-selective regions. Estimates of pRF size obtained with inverted face stimuli show similar patterns of variation (Figure S3).

#### pRF number

We next examined variations in the number of pRFs in each region (after filtering by *R^2^*, beta, and sigma thresholds). As in prior work, the number of pRFs decreased with eccentricity in all visual areas (Figure 5). The number of pRFs also decreased moving up the hierarchy, with lower numbers evident in face-selective areas and more dramatic reductions in pRF number in the periphery, consistent with a magnified foveal bias in these regions compared to early visual cortex.

**Figure 5.**
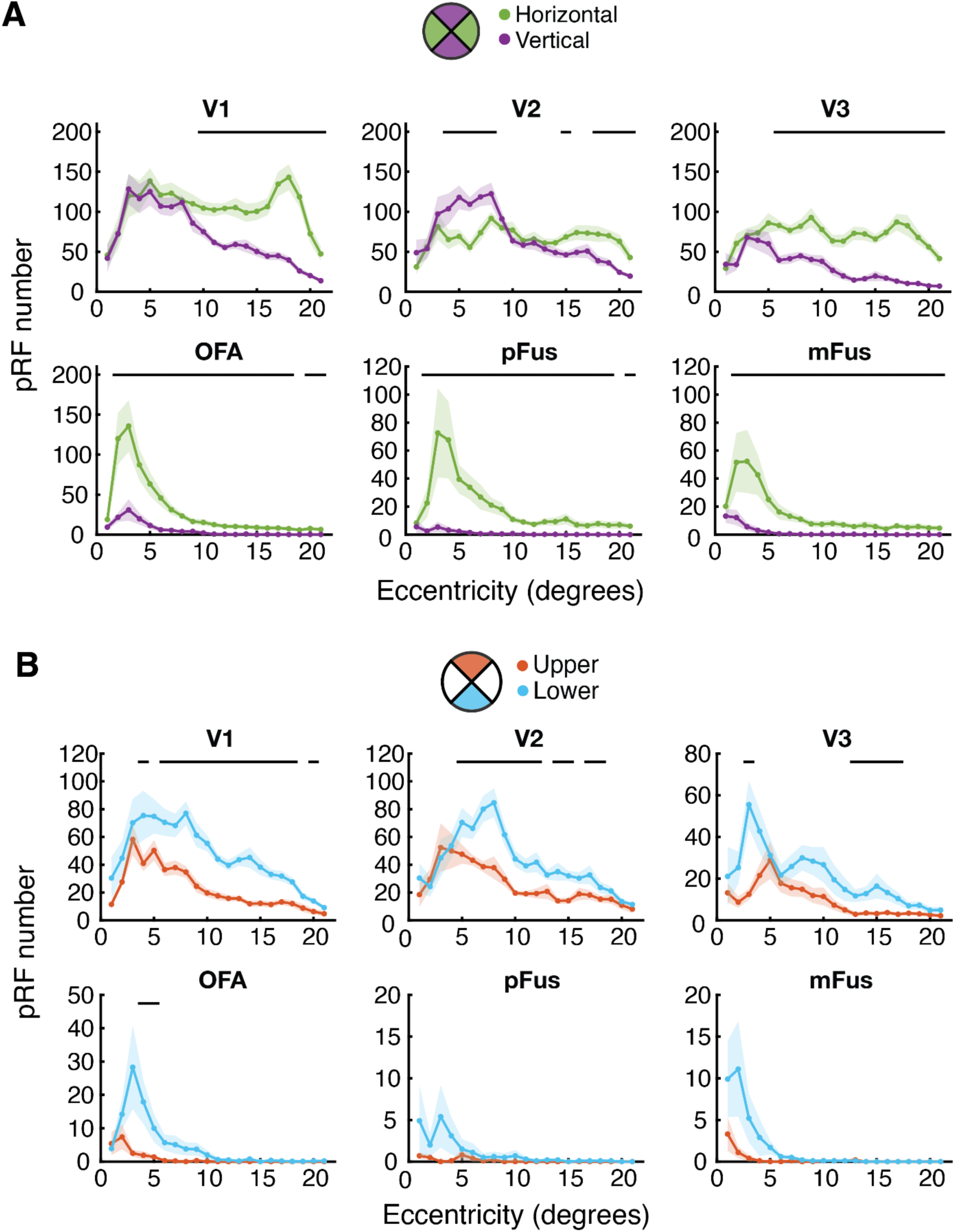
The mean number of pRFs responsive to upright face bars, plotted as a function of eccentricity along the horizontal (green) and vertical (purple) meridians **(A)** and in the upper and lower visual field **(B)**, plotted separately for early (V1-V3) and face-selective (OFA, pFus, mFus) cortical regions. Black lines indicate significant differences at each location (*p* < .05). Note differences in the y-axis scale across regions.

Figure 5A plots pRF numbers along the horizontal and vertical meridians, which show clear differences in early visual cortex. The main effect of location was significant in V1 (*F*_1,180_ = 26.83, *p* < .001) and V3 (*F*_1,180_ = 224.16, *p* < .001) driven by the greater number of pRFs on the horizontal meridian. Although there was no main effect of location in V2 (*F*_1,180_ = 0.01, *p* = .957), all three areas show a clear drop in pRF numbers on the vertical meridian with eccentricity, increasing the horizontal-vertical difference. Significant interactions between location and eccentricity were indeed evident in each case (V1: *F*_20,180_ = 14.64, *p* < .001; V2: *F*_20,180_ = 16.54, *p* < .001, V3; *F*_20,180_ = 7.94, *p* < .001). Wilcoxon tests showed that all three early visual areas had significantly more pRFs along the horizontal than vertical meridian in peripheral vision, with significant differences over a greater proportion of the visual field in V3 compared to V1 and V2. In face-selective cortex, a strong foveal bias was evident in all three areas, with pRF numbers dropping markedly at eccentricities beyond 5-10°. All three face-selective regions nonetheless show greater pRF numbers along the horizontal than vertical meridian, confirmed by main effects of location in each (OFA: *F*_1,180_ = 16.49, *p* = .003; pFus: *F*_1,160_ = 15.92, *p* = .004, mFus; *F*_1,180_ = 7.27, *p* = .025). There were again interactions between location and eccentricity in all three regions (OFA: *F*_20,180_ = 15.86, *p* < .001; pFus: *F*_20,160_ = 7.81, *p* < .001, mFus; *F*_20,180_ = 6.53, *p* < .001), though here Wilcoxon tests reveal significant differences both near the fovea and in the periphery. This over-representation of the horizontal meridian follows the predicted direction for behavioural anisotropies in both early and face-selective cortex.

Figure 5B plots differences in the lower vs. upper field, where main effects of location in V1-V3 confirmed a greater number of pRFs in the lower field, as predicted by behavioural anisotropies (V1: *F*_1,180_ = 36.19, *p* < .001; V2: *F*_1,180_ = 34.59, *p* < .001, V3; *F*_1,180_ = 10.37, *p* = .011). Interactions between location and eccentricity were also significant (V1: *F*_20,180_ = 3.86, *p* < .001; V2: *F*_20,180_ = 7.45, *p* < .001, V3; *F*_20,180_ = 2.68, *p* < .001), with Wilcoxon tests showing that the upper-lower difference generally increased towards the periphery. Upper-lower differences in pRF number were even more pronounced in face-selective regions, with all three face-selective areas showing strikingly few pRFs in the upper field. This gave main effects of location in OFA (*F*_1,180_ = 7.87, *p* = .021) and mFus (*F*_1,180_ = 19.06, *p* = .002), with more pRFs in the lower than upper field, consistent with predictions. Although there was no main effect of location in pFus (*F*_1,160_ = 2.93, *p* = .125), the interaction between location and eccentricity was significant for all three areas (OFA: *F*_20,180_ = 5.05, *p* < .001; pFus: *F*_20,160_ = 2.92, *p* < .001; and mFus: *F*_20,180_ = 11.29, *p* < .001). Wilcoxon tests showed that upper-lower differences were more pronounced near the fovea in OFA. Although the Wilcoxon tests did not find significant upper-lower differences in pFus and mFus, this is likely related to the low number of pRFs along the vertical meridian in these regions. Estimates of pRF number obtained with inverted face stimuli show similar patterns (Figure S4). Together, we observe more pRFs along the horizontal vs. vertical meridian and in the lower vs. upper field across both early and face-selective brain regions, in a manner consistent with behavioural anisotropies.

#### Visual-field coverage

Although the above variations in pRF size and number are somewhat mixed, we can examine their joint operation by computing pRF coverage around the visual field, as shown in Figure 3. Each brain region shows some degree of coverage throughout the periphery, though there is a clear decline with eccentricity, most rapidly in face-selective areas. To examine variations in coverage around the visual field, we next separated these coverage values by visual field location (Figure 6). Comparing the horizontal and vertical meridians in early visual cortex (Figure 6A), there were significant main effects of location in all three areas (V1: *F*_1,180_ = 8.36, *p* = .018; V2: *F*_1,180_ = 9.32, *p* = .014; V3: *F*_1,180_ = 46.57, *p* < .001), driven by higher coverage along the horizontal compared to the vertical meridian, consistent with the pattern of behavioural variations. There were also significant interactions between location and eccentricity in all three regions (V1: *F*_20,180_ = 30.78, *p* < .001; V2: *F*_20,180_ = 31.70, *p* < .001; V3: *F*_20,180_ = 59.97, *p* < .001), given that horizontal-vertical differences were most evident at intermediate eccentricities. Across early visual cortex, t-tests revealed differences at the majority of eccentricities tested, with the horizontal-vertical anisotropy becoming particularly pronounced in V3 relative to the earlier areas.

**Figure 6.**
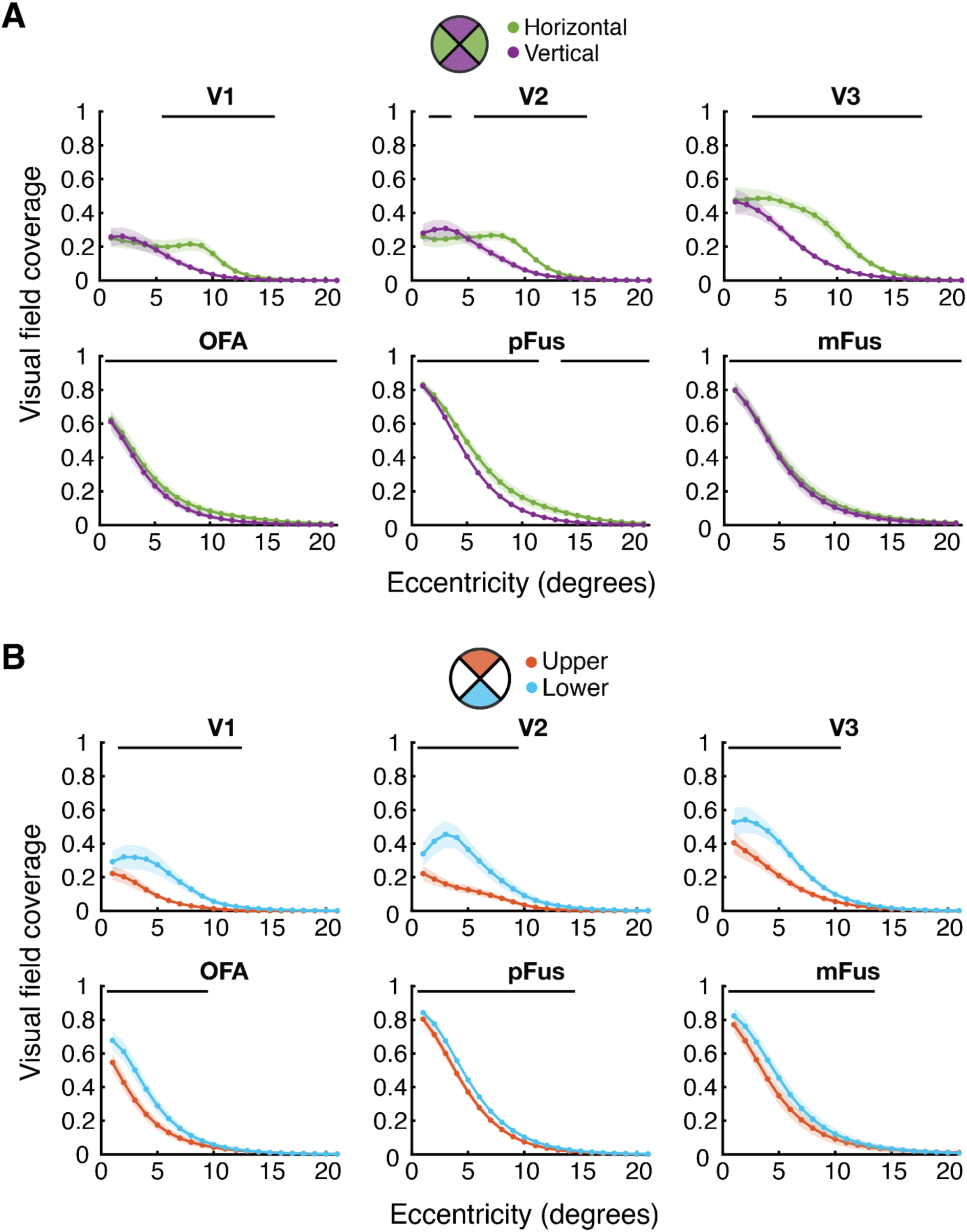
Mean visual-field coverage across eccentricity along the horizontal (green) and vertical (purple) meridians **(A)** and in the upper and lower visual field **(B)**, plotted separately for early (V1-V3) and face-selective (OFA, pFus, mFus) cortical regions. Black lines indicate significant differences at a given location (*p* < .05).

Face-selective brain regions all showed small but consistent horizontal-vertical anisotropies in visual-field coverage, again with greater coverage on the horizontal meridian and significant main effects of location in all three areas (OFA: *F*_1,180_ = 11.84, *p* = .007; pFus: *F*_1,160_ = 7.20, *p* = .028; mFus: *F*_1,180_ = 14.94, *p* = .004). There were also interactions between location and eccentricity in OFA (*F*_20,180_ = 3.17, *p* < .001) and pFus (*F*_20,160_ = 8.67, *p* < .001) but not in mFus (*F*_20,180_ = 0.27, *p* = .999), with t-tests confirming that these horizontal-vertical differences were present across most eccentricities in pFus, and all of them in OFA and mFus (the latter being small but nonetheless consistent). Altogether, these results reveal a consistent horizontal-vertical difference across the visual field within face-selective cortex, in a direction consistent with behavioural anisotropies.

Coverage also differed in the upper vs. lower field throughout V1-V3, with greater coverage in the lower vs. upper field, again consistent with behavioural effects. The main effect of location was accordingly significant in all areas (V1: *F*_1,180_ = 25.67, *p* < .001; V2: *F*_1,180_ = 31.87, *p* < .001; V3: *F*_1,180_ = 19.72, *p* = .002). All three regions showed significant interactions between location and eccentricity (V1: *F*_20,180_ = 30.79, *p* < .001; V2: *F*_20,180_ = 51.05, *p* < .001; V3: *F*_20,180_ = 32.37, *p* < .001), with t-tests confirming significant upper-lower differences only for eccentricities below 10°. Face-selective regions also had greater visual-field coverage in the lower vs. upper field, shown by main effects of location in all three areas (OFA: *F*_1,180_ = 16.09, *p* = .003; pFus: *F*_1,160_ = 10.32, *p* = .012; mFus: *F*_1,180_ = 24.80, *p* < .001). Like early visual cortex, there were also interactions between location and eccentricity (OFA: *F*_20,180_ = 41.28, *p* < .001; pFus: *F*_20,160_ = 13.21, *p* < .001; mFus: *F*_20,180_ = 18.49, *p* < .001), with t-tests again revealing significant differences at eccentricities below 10-15° but not beyond. Estimates of visual-field coverage obtained using inverted face stimuli show similar patterns (Figure S5).

Overall, coverage was consistently higher along the horizontal than vertical meridian and in the lower than upper field in both early visual cortex and face-selective areas. These variations in coverage, along with those for pRF number, demonstrate that low– and high-level visual areas show the same anisotropies in visual field sampling. This commonality does not however extend to pRF size, which showed more variable patterns that are unable to account for the pattern of behavioural anisotropies for face recognition.

### Inversion

Given that the spatial properties of both early and face-selective cortex vary in a manner consistent with behavioural anisotropies in face recognition, these properties may also vary with face orientation, following suggestions that smaller pRFs drive the advantage for upright over inverted face recognition (Poltoratski et al., 2021). During scans, to ensure that we engaged these face-selective processes, participants performed a gender recognition task. Performance showed a clear advantage for upright faces (Figure S6A), replicating the well-established inversion effect (Yin, 1969; Rossion and Gauthier, 2002) and demonstrating that our task was sufficient to engage configural face-recognition processes. This difference in difficulty did not affect fixation, with performance on the concurrent colour-change task at fixation equivalent with upright and inverted faces (Figure S6B).

pRF properties were examined for upright vs. inverted faces, here regardless of visual field location (pooling across the whole field). These properties did not differ in early visual cortex, as one would expect given the lack of selectivity for face orientation (shown in Figure S7). We focus instead on face-selective regions. Previous work has reported larger pRFs for upright compared to inverted faces in mFus and pFus but not OFA (Poltoratski et al., 2021). Here we observe larger pRFs for *inverted* than upright faces in OFA (Figure 7A), with a significant main effect of inversion (*β* = 0.97 [0.32, 1.61], *p* = .003). In pFus, pRF sizes were similar in both upright and inverted conditions, with a non-significant main effect (*β* = 0.32 [-0.46, 1.11], *p* = .416). This was also the case in mFus – despite a trend towards larger pRFs in the periphery for upright faces, the main effect of inversion was non-significant (*β* = –0.58 [-2.45, 1.28], *p* = .538). Wilcoxon tests further showed only sporadic significant differences with inversion at specific eccentricities within the three areas. As with the location variations discussed above, we conclude that pRF size was not modulated by inversion in a consistent manner.

**Figure 7.**
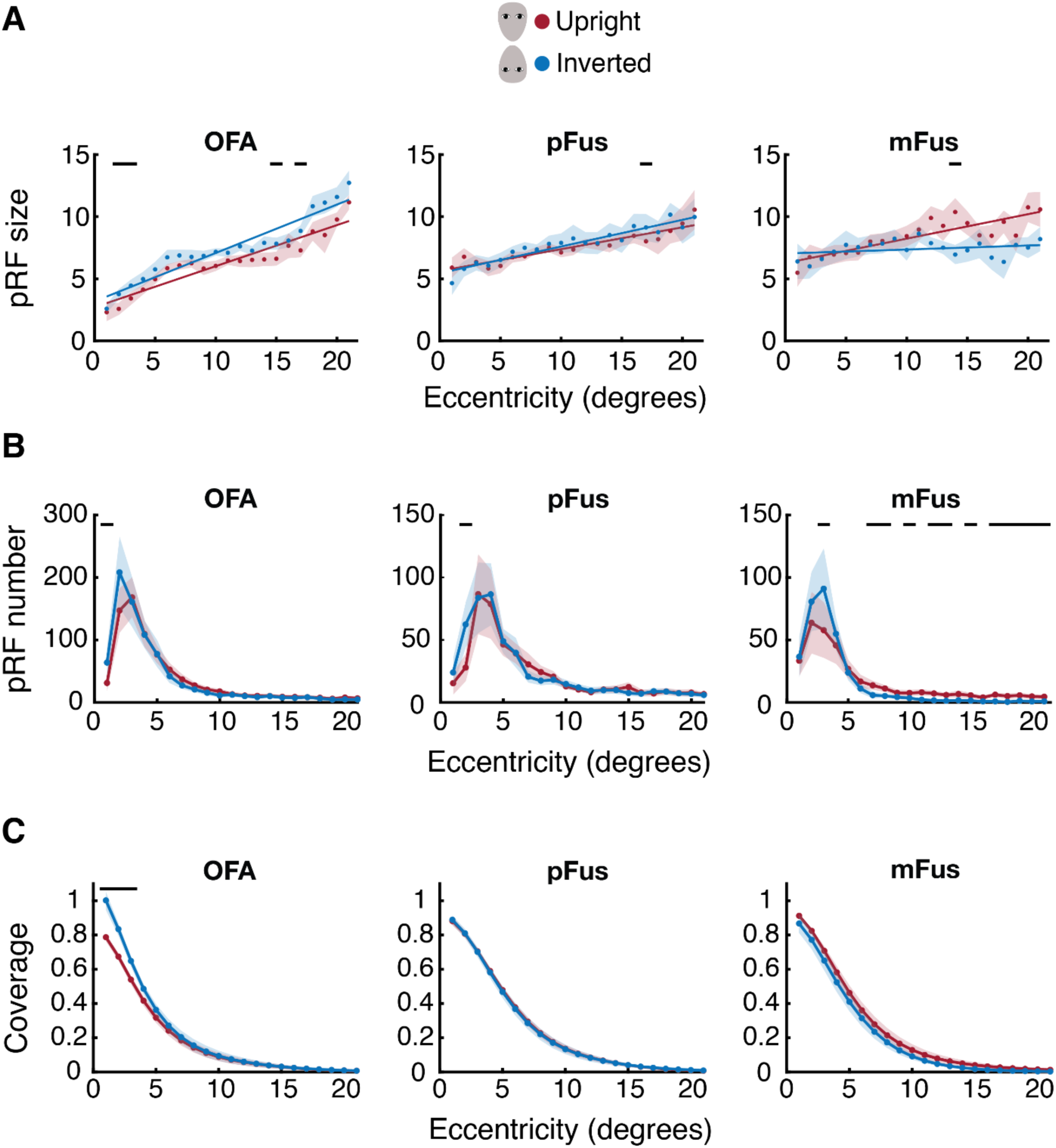
The effect of face orientation on spatial selectivity in face-selective regions, plotting mean pRF size **(A)**, pRF number **(B)** and visual-field coverage **(C)** across eccentricity for upright (red) and inverted (blue) faces, separately in the OFA, pFus and mFus. For each property, black lines indicate significant differences with face inversion (*p* < .05).

For pRF number (Figure 7B), there were no main effects of inversion in any of the face-selective regions (OFA: *F*_1,180_ = 0.24, *p* = .634; pFus: *F*_1,160_ = 2.48, *p* = .154; mFus: *F*_1,180_ = 0.53, *p* = .484). However, there were significant interactions between inversion and eccentricity in all three areas (OFA: *F*_20,180_ = 2.69, *p* < .001; pFus: *F*_20,160_ = 1.97, *p* = .011; mFus: *F*_1,180_ = 1.72, *p* = .034). Wilcoxon tests did not find consistent upright-inverted differences in OFA and pFus, though in mFus there were significantly more pRFs for upright than inverted faces over the majority of eccentricities, particularly beyond the fovea (despite one eccentricity near the fovea with more pRFs for inverted faces). This variation in the number of pRFs in mFus thus largely follows a pattern that could drive the behavioural Face Inversion Effect.

Finally, estimates of visual-field coverage (Figure 7C) were higher near the fovea for inverted vs. upright faces in OFA, though the main effect of inversion was non-significant (*F*_1,180_ = 0.02, *p* = .884), as was the interaction between inversion and eccentricity (*F*_20,180_ = 0.97, *p* = .496). These effects were similarly non-significant in pFus, both for the main effect (*F*_1,160_ = 0.13, *p* = .731) and interaction (*F*_20,160_ = 0.31, *p* = .998). Although the pattern seen in OFA was reversed in mFus, with slightly higher coverage levels for upright vs. inverted faces, the main effect (*F*_1,180_ = 2.18, *p* = .174) and interaction with eccentricity (*F*_20,180_ = 1.00, *p* = .460) were both non-significant. The lack of variation in coverage with face inversion suggests that this property could not drive differences in performance with upright vs. inverted faces, similar to the observation above that that pRF sizes do not differ reliably across these conditions. Variations in the number of pRFs sensitive did nonetheless follow the expected pattern in mFus, with greater numbers of pRFs found to be responsive to upright vs. inverted faces.

## Discussion

We show that the spatial selectivity of face-selective cortex (including OFA, pFus, and mFus) follows the same anisotropic pattern around the visual field as earlier retinotopic areas V1-V3. All of these areas show a greater *number* of population receptive fields (pRFs) and higher *visual-field coverage* along the horizontal than the vertical meridian and in the lower than upper field. These common variations suggest a continuity of visuospatial coding in the visual system, where high-level category-selective cortex encodes objects using the same spatial framework established in early regions (Groen et al., 2022). This pattern is consistent with behavioural anisotropies in both low-level vision (Carrasco et al., 2001; Abrams et al., 2012; Greenwood et al., 2017; Barbot et al., 2021; Himmelberg et al., 2023a) and higher-level face perception (Roux-Sibilon et al., 2023; Morsi et al., 2024). In contrast, *pRF sizes* did not vary in a consistent manner around the visual field in any of these regions, suggesting this property is poorly suited to explain these behavioural effects. Similarly, neither pRF sizes nor coverage varied with the upright vs inverted orientation of faces, contrary to prior reports (Poltoratski et al., 2021). We did however observe a reduction in pRF number in mFus for inverted faces, suggesting that similar sampling principles underlie both the spatial and stimulus selectivity of face-selective regions.

In early visual cortex, our observation of greater pRF numbers and increased visual-field coverage along the horizontal than vertical meridian and in the lower than upper field is consistent with previous findings (Amano et al., 2009), as well as prior demonstrations of greater cortical magnification and surface area in these locations (Silva et al., 2018; Himmelberg et al., 2023b). We show that these variations also arise in face-selective regions, with OFA, pFus and mFus all showing the same variations in pRF number and coverage (though the upper-lower difference in pRF number did not reach significance in pFus). In each case, horizontal-vertical differences were more pronounced than upper-lower differences, consistent with behavioural studies showing that the upper-lower difference is smaller in both low-level vision (Greenwood et al., 2017; Barbot et al., 2021; Kurzawski et al., 2021) and face perception (Morsi et al., 2024).

As in prior studies, we observe a strong foveal bias in face-selective regions (Kay et al., 2015; Finzi et al., 2021; Poltoratski et al., 2021; Silson et al., 2022), with a steeper decline in pRF number with eccentricity than in early visual cortex. Despite these changes, the horizontal-vertical and upper-lower differences remain robust in these regions, suggesting that these anisotropies are actively maintained in category-selective cortex, rather than being passively inherited from early cortex. This active maintenance may reflect a sensitivity to the typical locations of faces in the visual field – for instance, over-sampling along the horizontal meridian may reflect an increased sensitivity to the plane along which we most typically encounter faces (e.g. given that groups of faces tend to appear at similar heights; de Haas et al., 2016). Similar principles are evident elsewhere in category-selective cortex, where the retinotopic properties of scene-selective areas show a reversed upper-lower anisotropy, matching the upper-field advantage in scene-recognition tasks (Silson et al., 2015).

Despite the sharp reductions in pRF numbers in peripheral vision, face-selective areas retained some degree of visual-field coverage remained across all eccentricities tested (up to ±21.65°). This level of visual-field coverage is sufficient to support the clear face-recognition abilities observed in peripheral vision (McKone, 2004; Kovacs et al., 2017; Kalpadakis-Smith et al., 2018; Roux-Sibilon et al., 2023; Morsi et al., 2024), as well as to detect and direct rapid saccades towards peripheral face stimuli (Crouzet et al., 2010; Boucart et al., 2016), albeit with some reduction in efficiency relative to foveal vision (Mäkelä et al., 2001). This peripheral coverage is driven by the large receptive fields in face-selective areas, which can thus be seen as an adaptive response to the reduced neural representation of the periphery.

Our results do not offer support for a common pattern of anisotropies in pRF size through the visual hierarchy. Though area V3 showed the expected pattern of larger pRFs along the vertical than horizontal meridian, these differences were not observed in V1 and V2. We also found *larger* pRFs in the lower than upper field across all three early visual areas, the opposite direction to that predicted by behavioural anisotropies. This is consistent with prior reports of little-to-no difference in pRF sizes along the vertical meridian (Himmelberg et al., 2023b), though it differs from reports of horizontal-vertical differences across V1-V3 (Silva et al., 2018) and smaller pRFs in the lower vs. upper field (Silson et al., 2018; Silva et al., 2018). These mixed findings may reflect the inherent variability in measurements of pRF size, which are known to be more susceptible to variations in stimulus properties or fitting procedures than properties like pRF location (Alvarez et al., 2015; van Dijk et al., 2016; Lage-Castellanos et al., 2020; Lerma-Usabiaga et al., 2021; Linhardt et al., 2021). Nonetheless, the lack of variation in pRF sizes for face-selective regions in our dataset consolidates these mixed findings in early cortex, suggesting that receptive field size is not the main factor driving differences in perception across the visual field.

The size of face-selective pRFs was also invariant to face orientation, contrary to prior results (Poltoratski et al., 2021). In addition to the above-noted challenges in measuring pRF size, these previously observed reductions could reflect their reduced beta amplitudes for inverted vs. upright faces, which can reduce goodness-of-fit (Schwarzkopf et al., 2014; Anderson et al., 2017). This in turn would bias averages towards smaller values since larger pRFs tend to drop out first (Alvarez et al., 2015; Hughes et al., 2019). A similar effect is unlikely to be present in our results, given similar beta amplitudes for upright and inverted faces (Figure S2C). We nonetheless found a clear behavioural inversion effect (Figure S6), confirming the engagement of face-specific processes during scans. More broadly, the invariance in pRF size with both inversion and visual field location does not support the proposed association between larger receptive fields and better face recognition (Witthoft et al., 2016; Avidan and Behrmann, 2021). Also inconsistent with this theory is our observation that pRF size increases with eccentricity within face-selective regions, as in prior studies (Kay et al., 2013; Poltoratski et al., 2021). If larger pRFs improved recognition, performance should improve in peripheral vision, contrary to established findings (Mäkelä et al., 2001; McKone, 2004). Together these findings suggest that the inversion effect cannot parsimoniously be explained by changes in pRF size.

We do however observe a greater number of pRFs for upright than inverted faces in area mFus, particularly in the periphery. Changes in the number of face-selective neurons with inversion could therefore drive the differences in these abilities, consistent with the reduced BOLD signal with inversion in face-selective regions (Kanwisher et al., 1998). The greater number of responsive voxels for upright faces may reflect the recruitment of neurons selective for the configural properties and/or natural statistics of upright faces (Le Grand et al., 2001; Rossion, 2008). This mirrors the variation in pRF numbers around the visual field, where more pRFs were associated with better performance, suggesting a common principle whereby greater sampling resources produce higher sensitivity. However, the inversion-related differences in pRF number were smaller than the variations across the visual field, despite the robust behavioural inversion effect and its consistency in prior work. The inversion effect may also therefore reflect qualitative differences in the firing of face-selective neurons, rather than purely quantitative differences in the number of active neurons.

Altogether our results demonstrate that face-selective cortex samples the visual field similarly to early visual cortex, with greater pRF numbers and visual-field coverage along the horizontal than vertical meridian and in the lower than upper field. These retinotopic variations align with the behavioural anisotropies observed for both low-level visual abilities (Carrasco et al., 2001; Abrams et al., 2012; Greenwood et al., 2017; Barbot et al., 2021; Himmelberg et al., 2023a) and face perception (Roux-Sibilon et al., 2023; Morsi et al., 2024). In contrast, pRF sizes did not reliably follow the same pattern as these behavioural anisotropies in either early or face-selective regions. Although large receptive fields are likely necessary to support face-recognition abilities in peripheral vision, given the pronounced foveal bias in face-selective regions, variations in size alone cannot explain performance differences around the visual field. We further observe that pRF numbers in area mFus decrease with face inversion, suggesting that the advantage for upright over inverted faces may at least in part reflect changes in the resources used in sampling the visual scene. This convergence suggests that variations in performance around the visual field and across upright vs. inverted faces are linked to a common principle: the allocation of neural sampling resources. More broadly, the common spatial selectivity in early and face-selective cortex is consistent with a unified visuospatial framework throughout the visual system (Groen et al., 2022).

## Acknowledgements

Funded by the Biotechnology and Biological Sciences Research Council [BB/J014567/1] and the Academy of Medical Sciences [NGRI/1087]. Thanks to Jolien Schuurmans for supplying stimuli used in the functional localiser for face-selective brain regions.

## Supplementary Information

**Figure S1.**
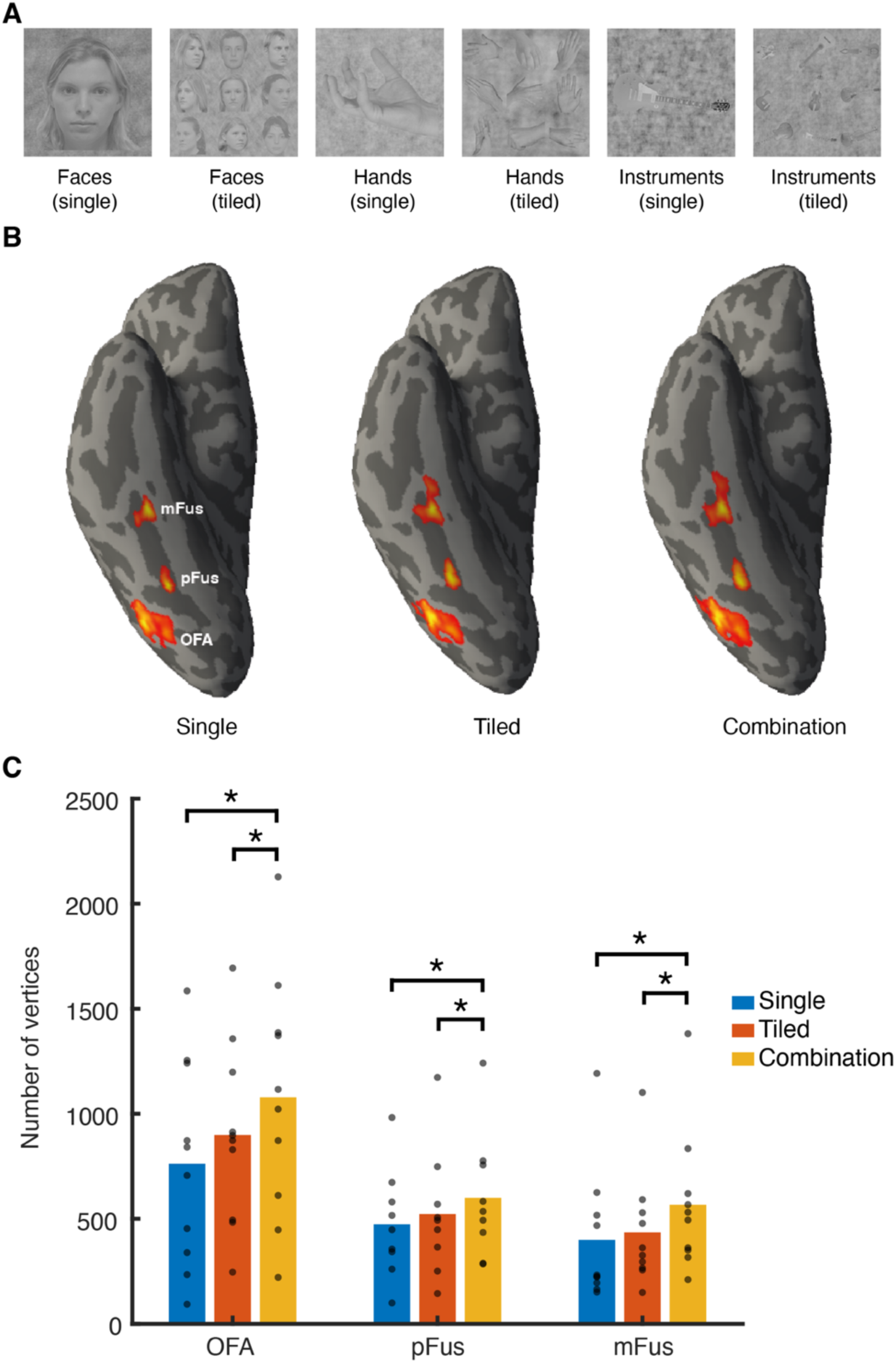
The localisation of face-selective brain regions. **A.** Stimuli used for the six image categories in the localiser runs. **B.** Face-selective regions OFA, pFus and mFus on a ventral view of an inflated model of the right cortical hemisphere in an example participant, defined using different localisation stimuli (single faces vs. single hands and instruments, tiled comparisons, or the combination of both single and tiled). In each case, ROIs have the same central location, with the activation spread out over a different amount of the cortical surface depending on the localisation approach. **C.** The number of vertices in each face-selective ROI, delineated using localiser stimuli consisting of single faces, tiled faces, or the combination of both. Dots show individuals and bars the mean number of vertices. Asterisks show significant differences (*p* < .05).

**Figure S2.**
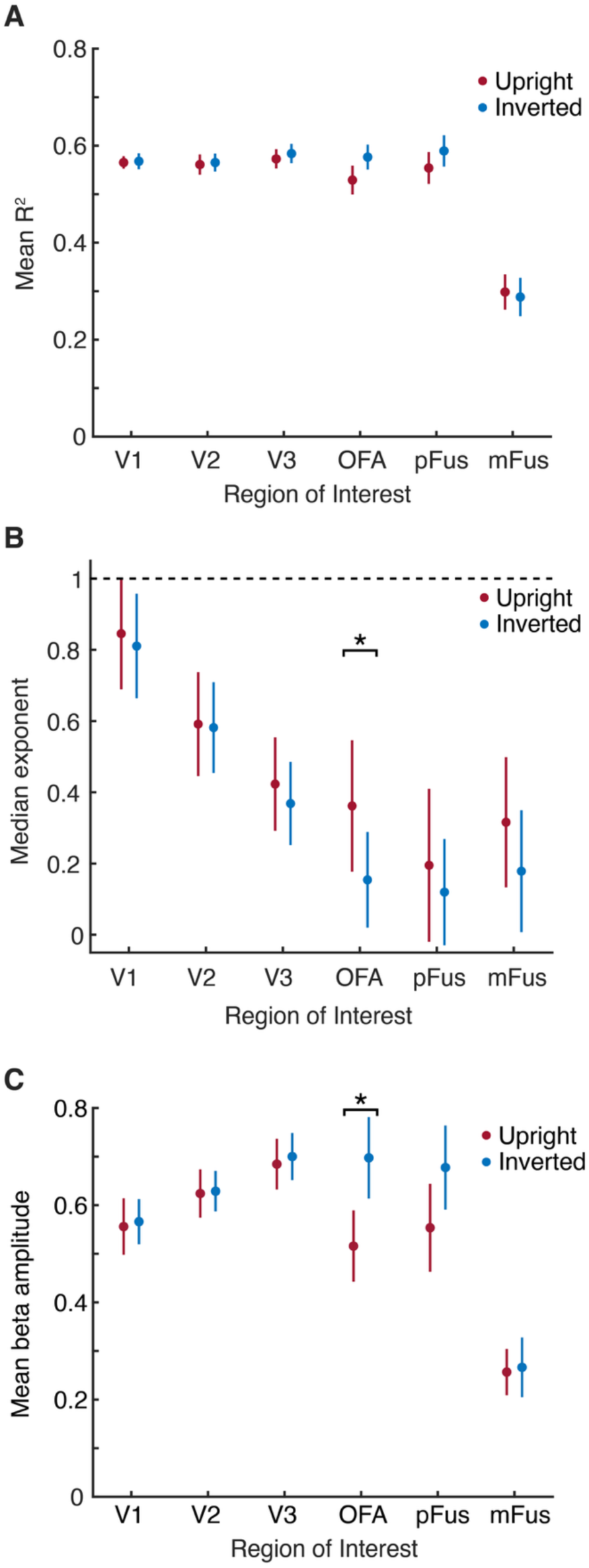
A. Mean R^2^ values for the compressive spatial summation (CSS) pRF model for upright (red) and inverted (blue) faces in each of the ROIs. Error bars show the SEM. **B.** Median exponent values for the CSS model in each ROI, plotted as in panel A. The dashed line represents linear summation, with all values < 1 indicating compression. Exponent values are lower (indicating increased compression) in face-selective regions compared to V1-V3, as in Kay et al. (2013). The asterisk denotes a statistically significant difference (*p* < .05). **C.** Mean beta amplitudes for the CSS model in each ROI, plotted as in other panels.

**Figure S3.**
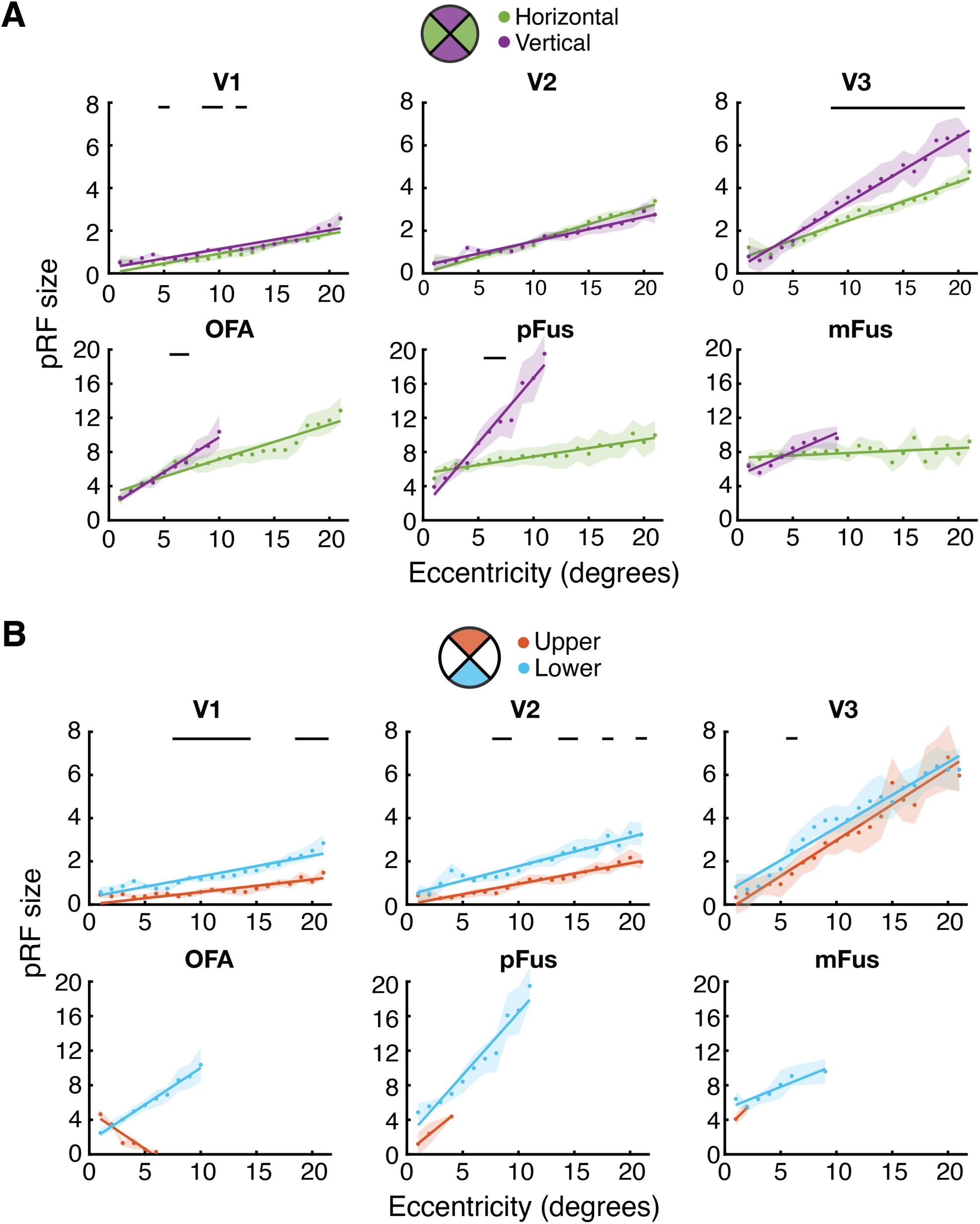
Mean pRF size across eccentricity measured with inverted faces, plotted along the horizontal (green) and vertical (purple) meridians **(A)** and in the upper and lower visual field **(B)**. At each eccentricity, size estimates were only plotted if they were averaged from at least five vertices. Black lines indicate significant differences according to location (*p* < .05). Note variations in the y-axis between regions.

**Figure S4.**
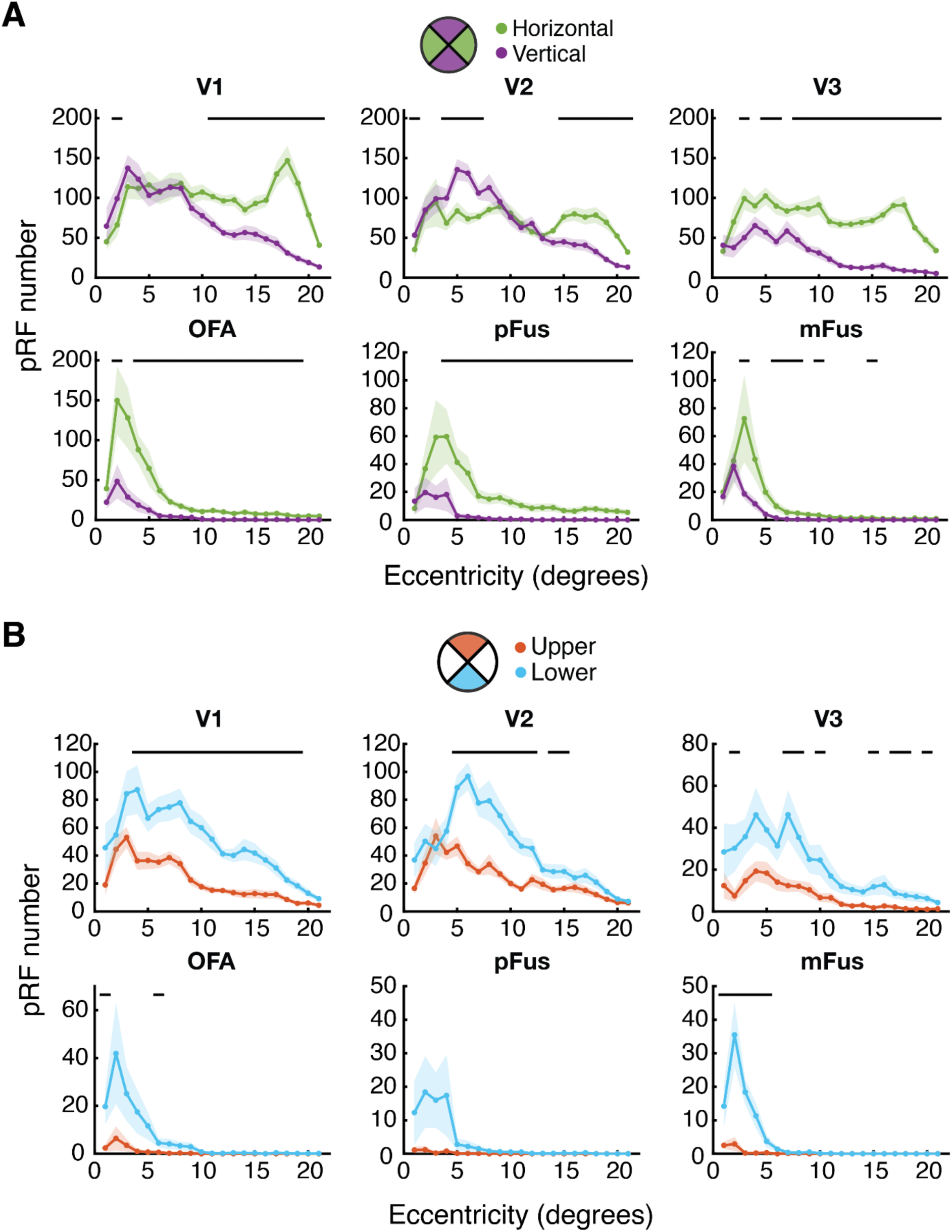
Mean pRF number for inverted faces, plotted across eccentricity on the horizontal (green) and vertical (purple) meridians **(A)** and in the upper and lower visual field **(B)**. Black lines indicate significant differences according to location (*p* < .05). Note variations in the y-axis between regions.

**Figure S5.**
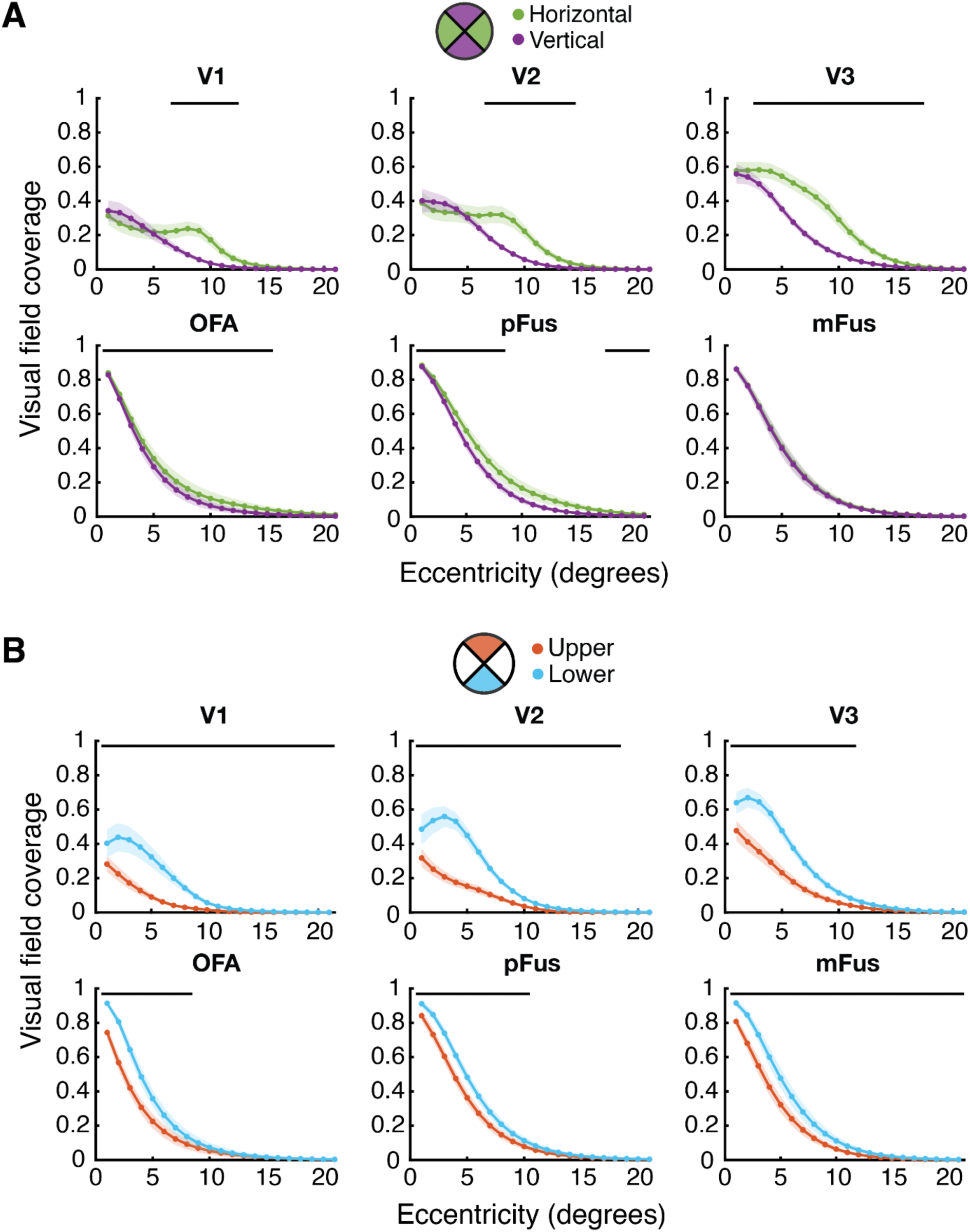
Visual-field coverage for inverted faces, plotted across eccentricity on the horizontal (green) and vertical (purple) meridians **(A)** and in the upper and lower visual field **(B)**. At each eccentricity, size estimates were only plotted if they were averaged from at least five vertices. Black lines indicate significant differences according to location (*p* < .05).

**Figure S6.**
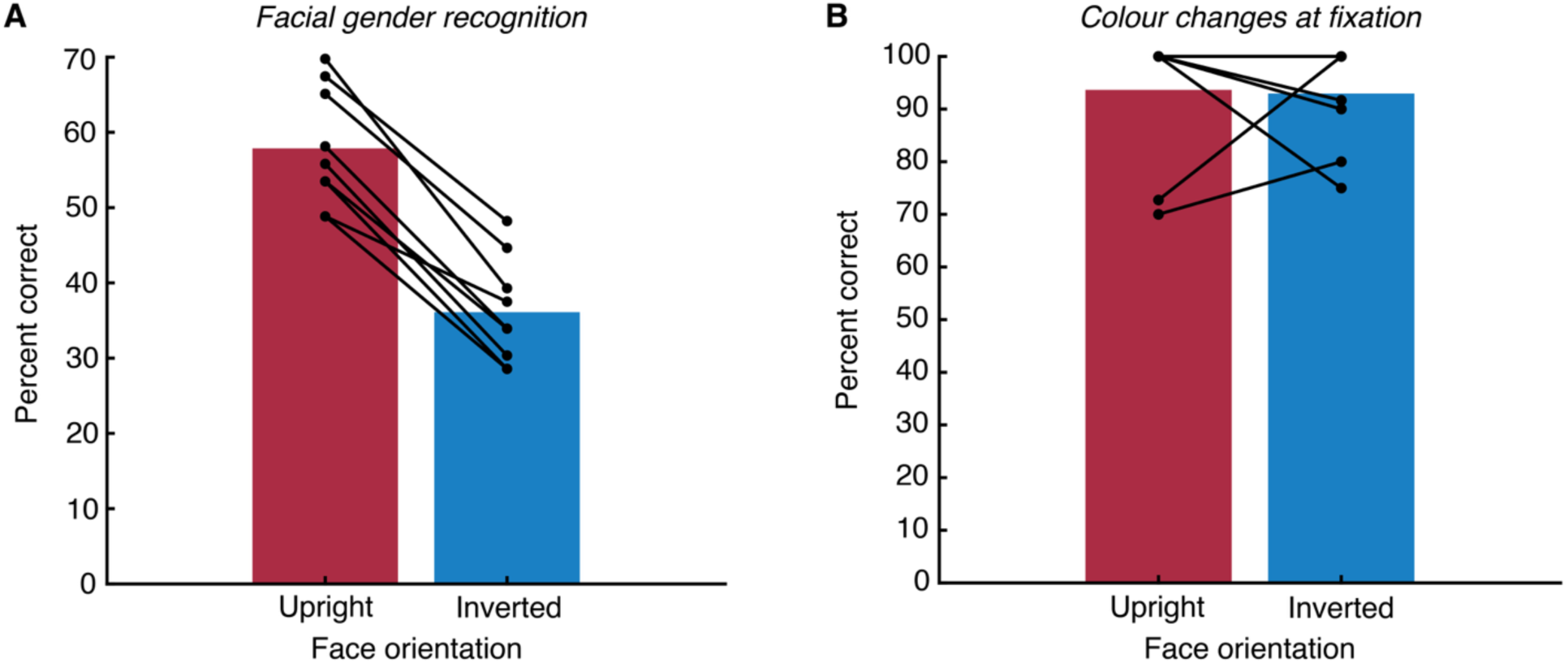
Behavioural results as a function of face orientation within the bar stimuli used to measure pRFs. **A.** Percent-correct recognition performance on the facial gender recognition task, showing the percentage of correctly identified male bars. Dots show individual data, with lines joining each participant’s performance for upright and inverted faces. A clear inversion effect is seen, with worst performance for inverted faces. **B.** Percent-correct recognition performance on the fixation task during the pRF experiment, showing the percentage of fixation cross colour changes that were correctly detected. Dots show individual data, with lines joining each participant’s performance for the fixation task during the presentation of upright and inverted faces. Four participants had 100% correct in both the upright and inverted runs. On the whole, there is no difference in performance when faces were upright vs. inverted.

**Figure S7.**
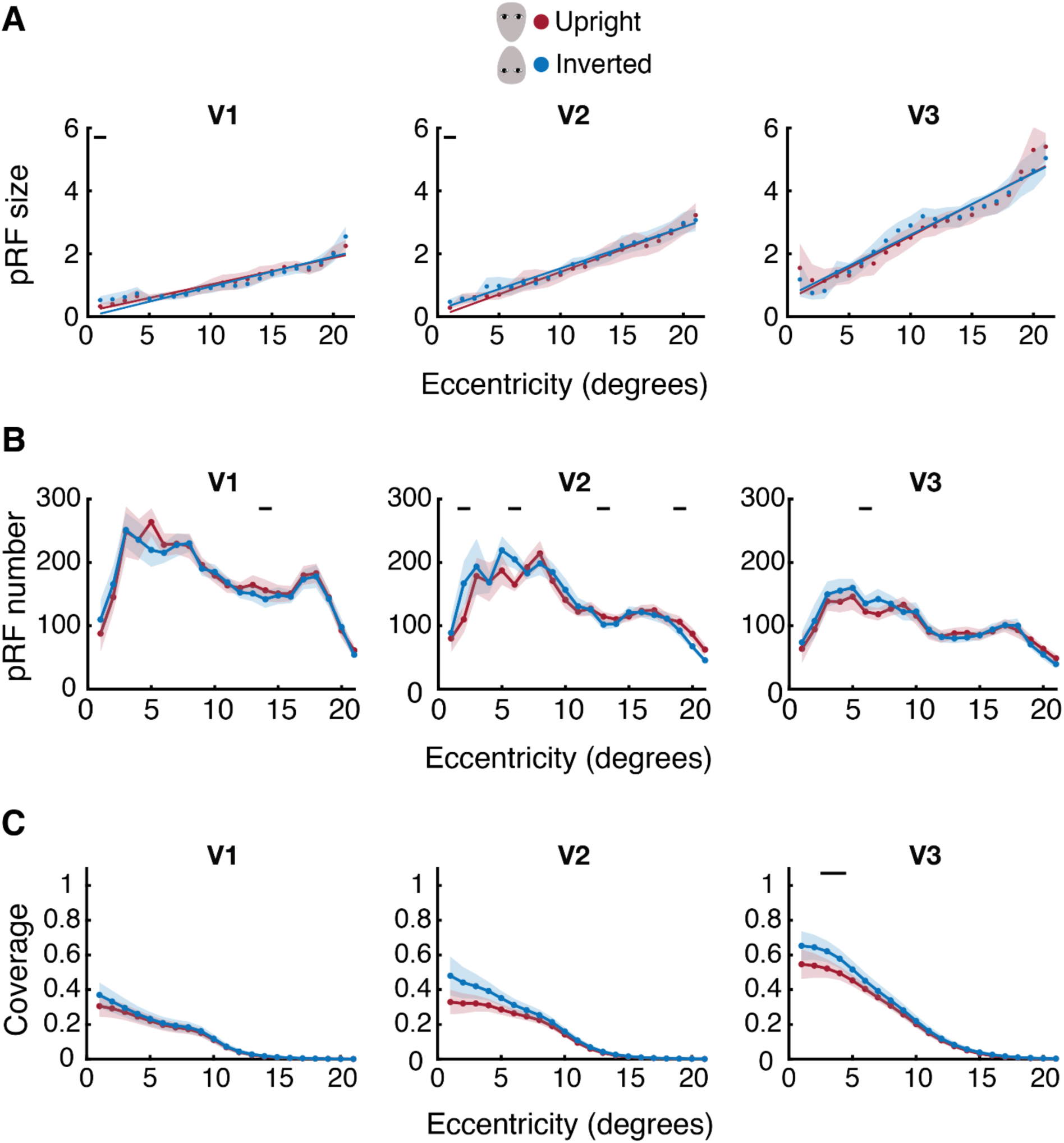
The effect of face orientation on spatial selectivity in early visual cortex. **A.** Mean pRF sizes in V1-V3, plotted across eccentricity for upright (red) and inverted (blue) faces. Black lines indicate significant differences in each property according to face inversion (*p* < .05). There were no significant main effects of inversion in any area (V1: *β* = –0.01 [-0.20, 0.18], *p* = .910; V2: *β* = 0.08 [-0.19, 0.34], *p* = .570; V3: *β* = 0.03 [-0.33, 0.40], *p* = .856). **B.** pRF number, plotted as in panel A. Main effects of inversion were again non-significant in all areas (V1: *F*_1,180_ = 0.32, *p* = .583; V2: *F*_1,180_ = 2.17, *p* = .175; V3: *F*_1,180_ = 1.45, *p* = .259). **C.** Visual-field coverage, plotted as in panel A. Main effects of inversion were again non-significant in all areas (all F<1).

