## Supplementary Information for "Face-selective cortical regions inherit the visuospatial organisation of early visual cortex"

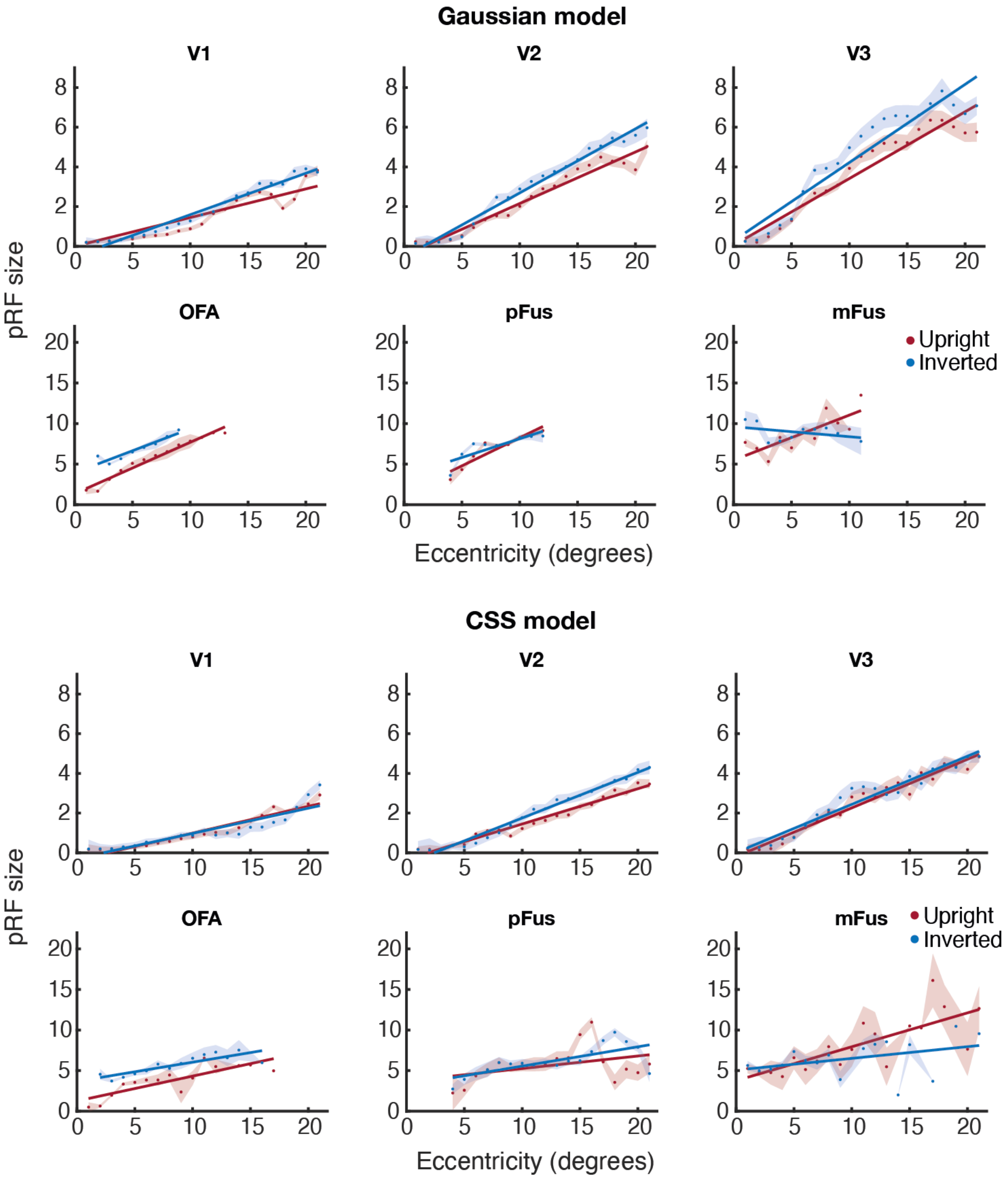


Figure S1. Mean estimates of pRF size in one participant, for the Gaussian (top) and compressive spatial summation (CSS; bottom) models, for upright (red) and inverted (blue) faces. The Gaussian model estimates pRF size as $\sigma$ (the standard deviation of the pRF, in degrees of visual angle), while the CSS model defines pRF size as $\sigma/\sqrt{(n)}$, with $n$ being the exponent of the compressive spatial summation parameter (Kay et al. 2013). Note the different axis scales between V1-V3 and the face-selective regions.

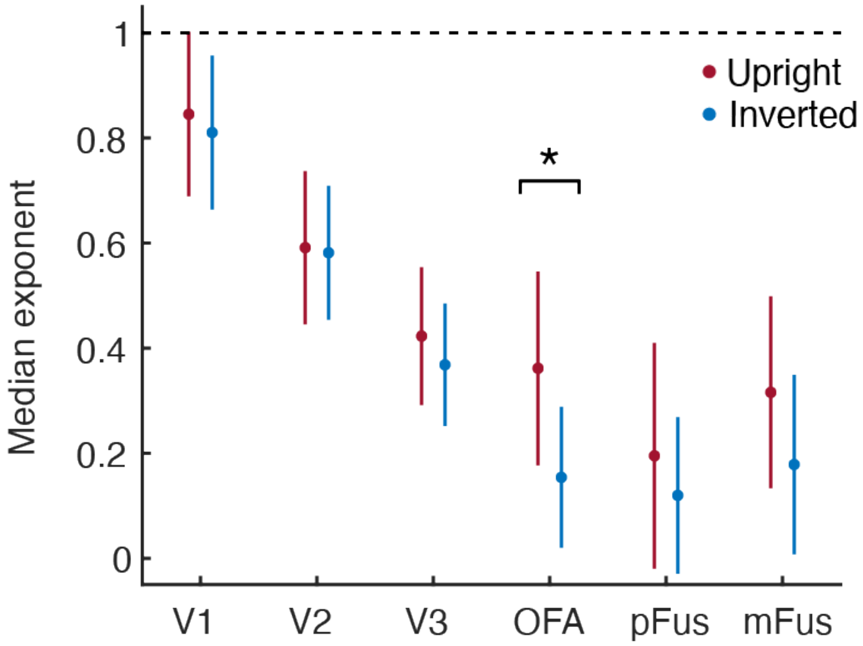


Figure S2. Median exponent values from the compressive spatial summation (CSS) pRF model for upright (red) and inverted (blue) faces in each of the ROIs. Error bars represent the SEM. The dashed line represents linear summation, with all values < 1 indicating compression. Exponent values are lower (indicating increased compression) in face-selective regions compared to V1-V3, as in Kay et al. (2013). The asterisk denotes a statistically significant difference ($p < .05$).

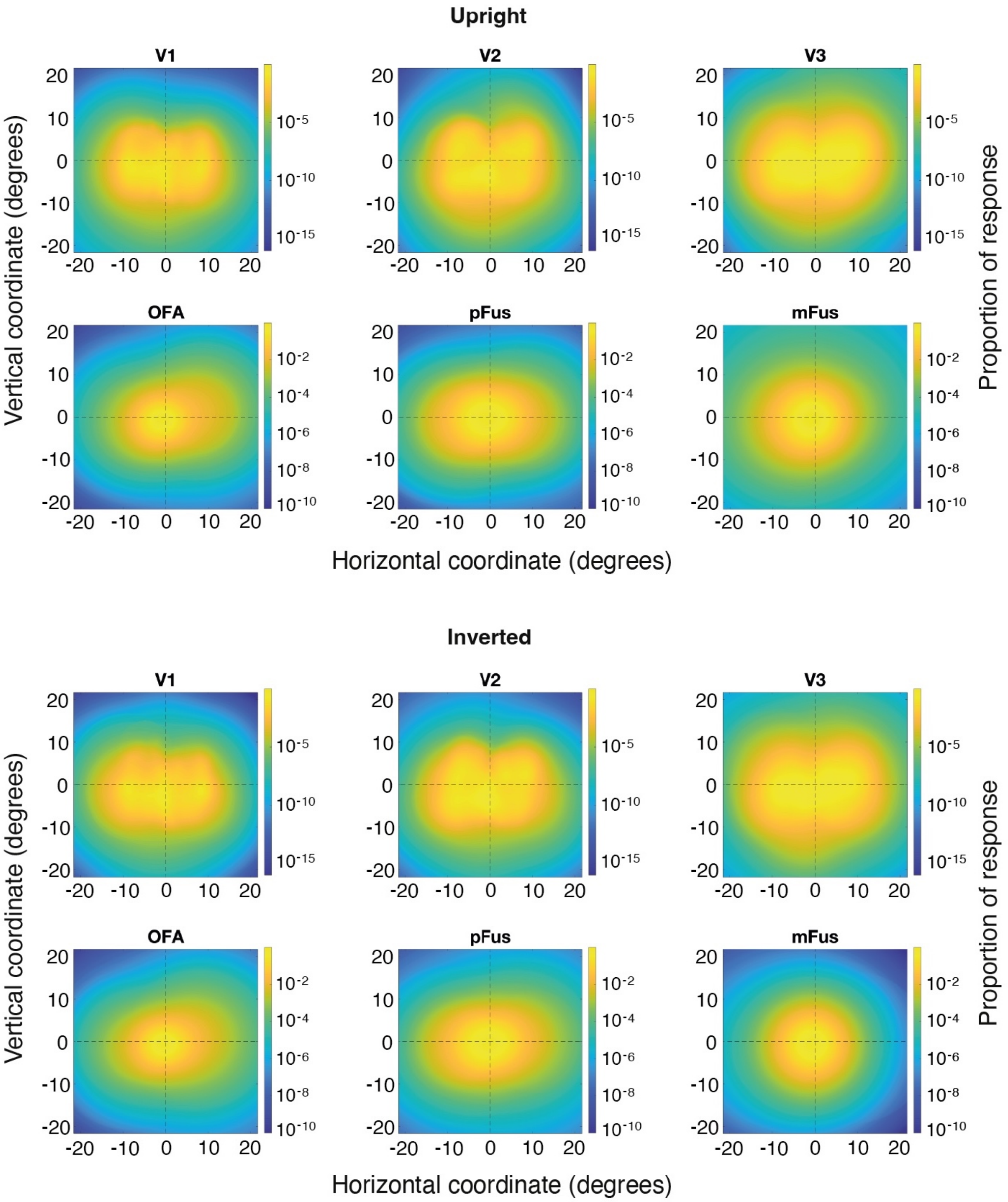


Figure S3. Mean visual field coverage for upright (top) and inverted (bottom) faces. Coordinates represent eccentricity (in degrees of visual angle), with negative values for the left and positive values for the right visual field. Values were converted to log scale before plotting, for visualisation purposes (see colour bar).

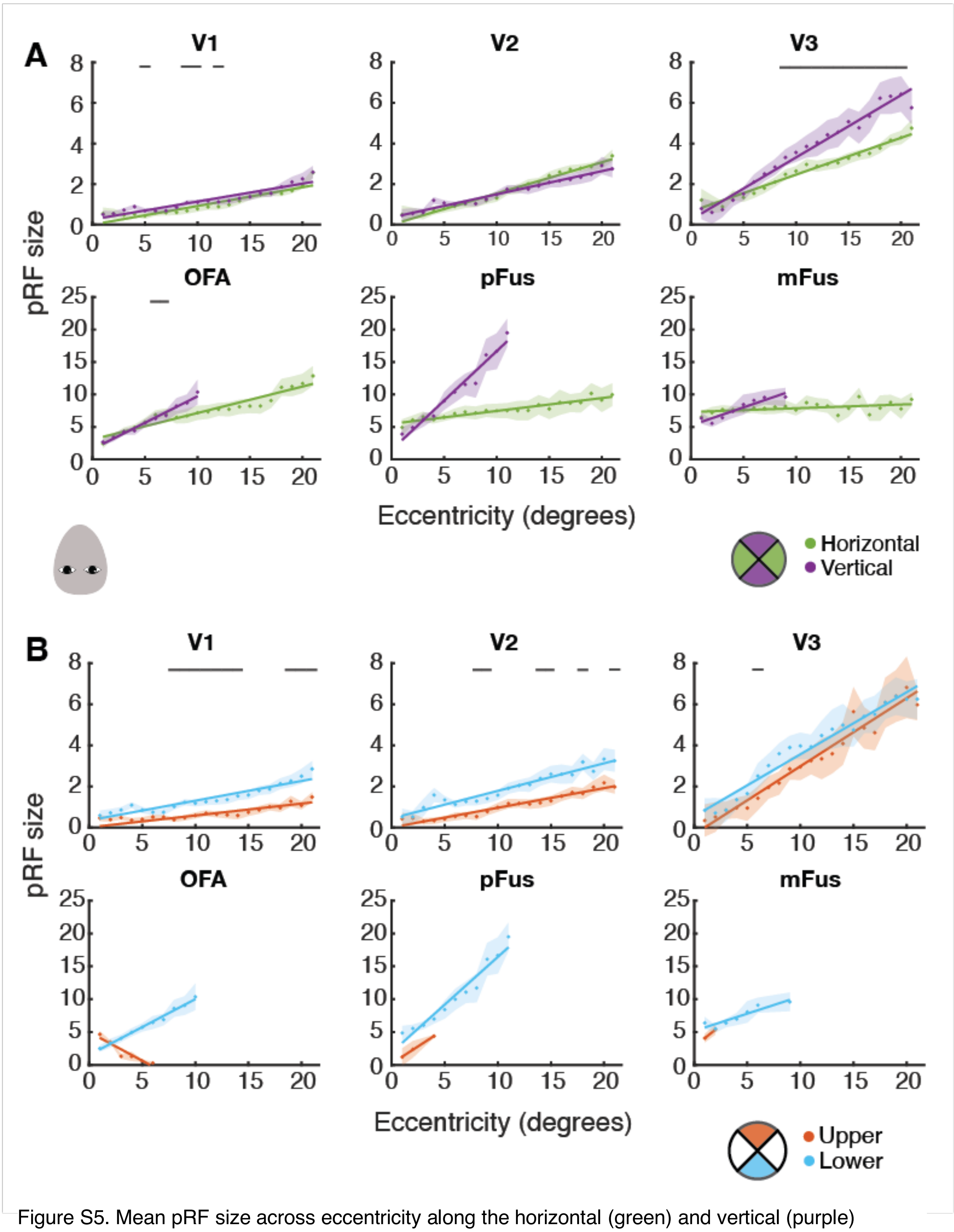


Figure S5. Mean pRF size across eccentricity along the horizontal (green) and vertical (purple) meridians **(A)** and in the upper and lower visual field **(B)** for the inverted faces. At each eccentricity, size estimates were only plotted if they were averaged from at least five vertices. Black lines indicate significant differences according to location ($p < .05$).

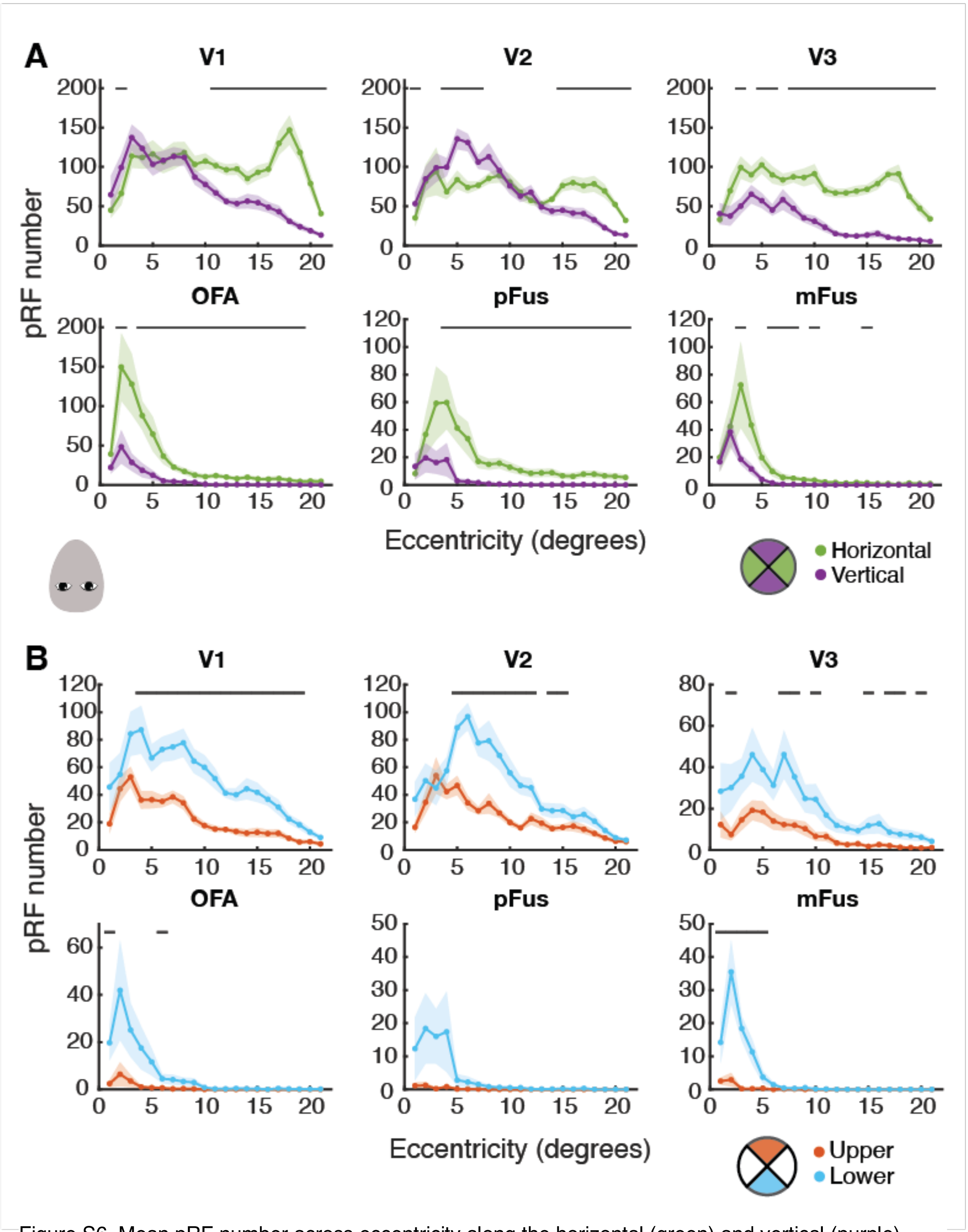


Figure S6. Mean pRF number across eccentricity along the horizontal (green) and vertical (purple) meridians **(A)** and in the upper and lower visual field **(B)** for the inverted faces. At each eccentricity, size estimates were only plotted if they were averaged from at least five vertices. Black lines indicate significant differences according to location ($p < .05$).

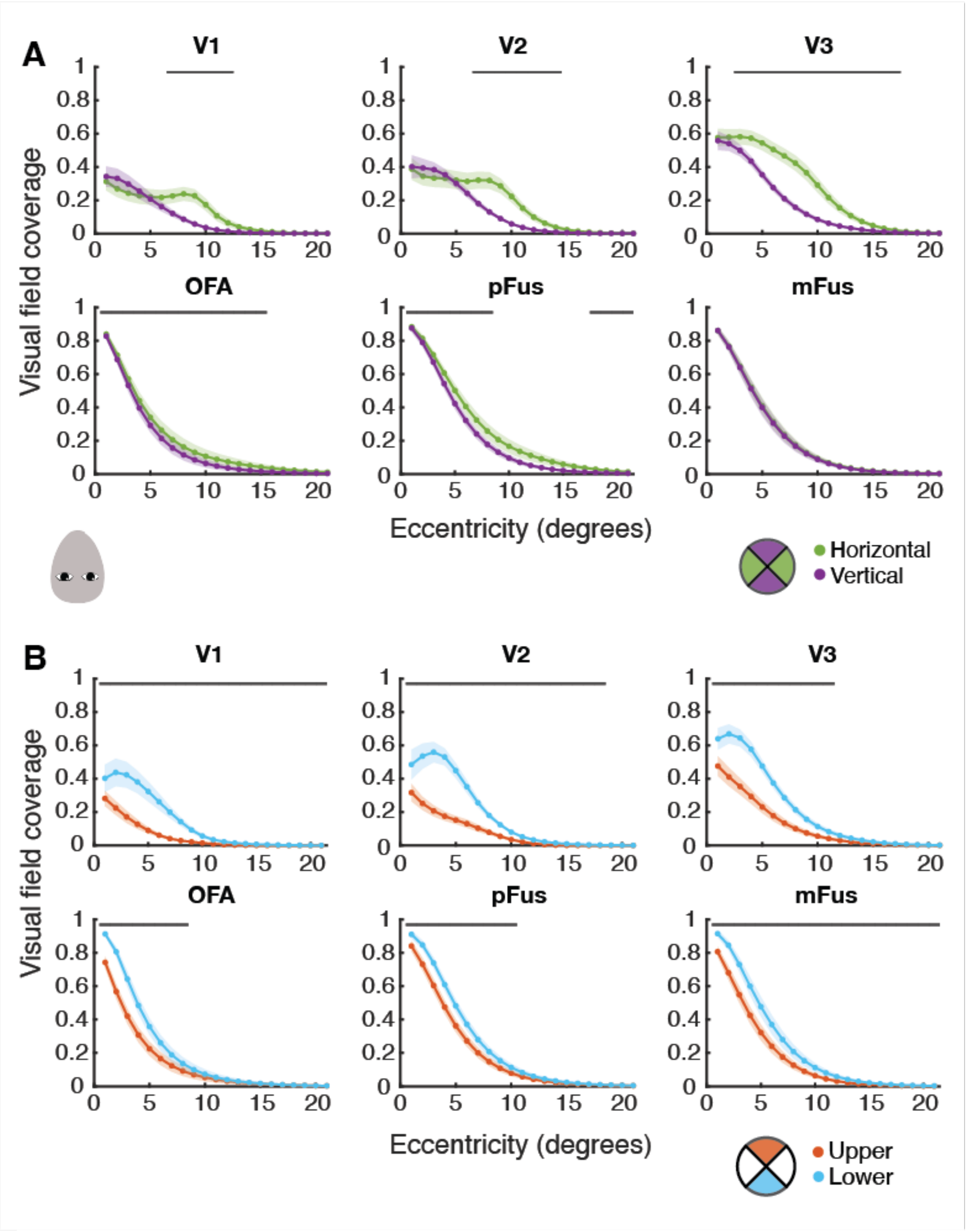


Figure S7. Visual field coverage across eccentricity along the horizontal (green) and vertical (purple) meridians **(A)** and in the upper and lower visual field **(B)** for the inverted faces. At each eccentricity, size estimates were only plotted if they were averaged from at least five vertices. Black lines indicate significant differences according to location ($p < .05$).

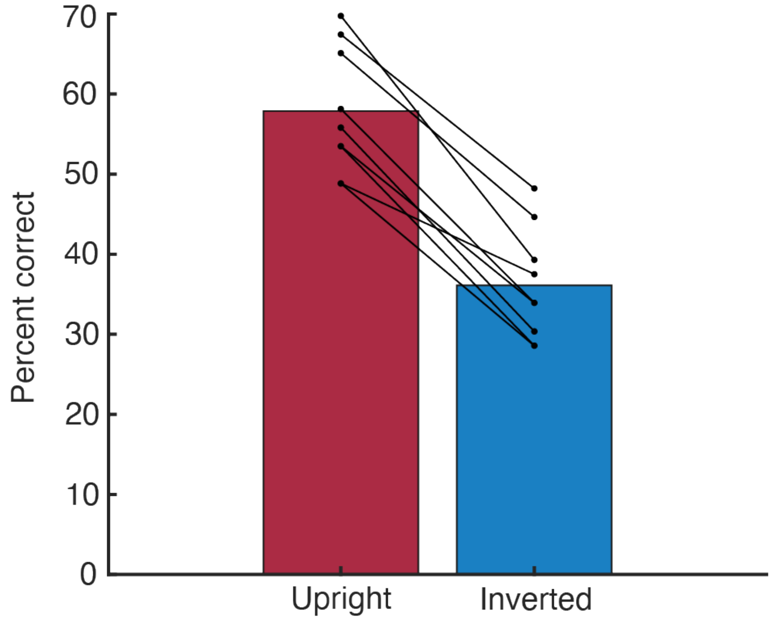


Figure S8. Behavioural results from the gender task in the pRF experiment, showing the percentage of correctly identified male bars. Dots show individual data, with lines joining each participant’s performance for upright and inverted faces.

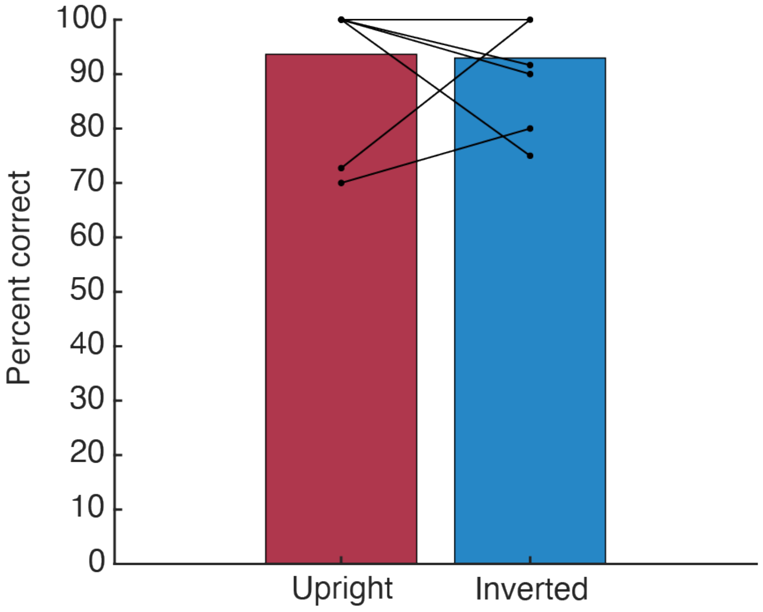


Figure S9. Behavioural results from the fixation task in the pRF experiment, showing the percentage of fixation cross colour changes that were correctly identified. Dots show individual data, with lines joining each participant's performance for upright and inverted faces. Four participants had 100% correct in both the upright and inverted runs.

| | ROI | Fixed factors | *β* | *p* | 95% CI |
|---|---|---|---|---|---|
| pRF size | V1 | Intercept | 0.22 | | |
| | | Inversion | -0.01 | .910 | -0.20, 0.18 |
| | | Eccentricity | 0.08 | **< .001** | 0.06, 0.10 |
| | V2 | Intercept | 0.05 | | |
| | | Inversion | 0.08 | .571 | -0.19, 0.34 |
| | | Eccentricity | 0.13 | **< .001** | 0.10, 0.16 |
| | V3 | Intercept | 0.53 | | |
| | | Inversion | 0.03 | .856 | -0.33, 0.40 |
| | | Eccentricity | 0.20 | **< .001** | 0.14, 0.26 |

| | ROI | Factors | *df* | *F* | *p* |
|---|---|---|---|---|---|
| pRF number | V1 | Inversion | 1, 180 | 0.32 | .583 |
| | | Eccentricity | 20, 180 | 13.67 | **< .001** |
| | | Inversion*Eccentricity | 20, 180 | 0.72 | .802 |
| | V2 | Inversion | 1, 180 | 2.17 | .175 |
| | | Eccentricity | 20, 180 | 10.37 | **< .001** |
| | | Inversion*Eccentricity | 20, 180 | 2.40 | **.001** |
| | V3 | Inversion | 1, 180 | 1.45 | .259 |
| | | Eccentricity | 20, 180 | 7.81 | **< .001** |
| | | Inversion*Eccentricity | 20, 180 | 0.67 | .852 |

| | ROI | Factors | *df* | *F* | *p* |
|---|---|---|---|---|---|
| Visual field coverage | V1 | Inversion | 1, 180 | 0.02 | .882 |
| | | Eccentricity | 20, 180 | 27.67 | **< .001** |
| | | Inversion*Eccentricity | 20, 180 | 11.18 | .965 |
| | V2 | Inversion | 1, 160 | 0.59 | .462 |
| | | Eccentricity | 20, 160 | 35.22 | **< .001** |
| | | Inversion*Eccentricity | 20, 160 | 1.77 | **.027** |
| | V3 | Inversion | 1, 180 | 0.03 | .862 |
| | | Eccentricity | 20, 180 | 76.84 | **< .001** |
| | | Inversion*Eccentricity | 20, 180 | 0.87 | .625 |

Figure S10. Linear mixed effects model (pRF size) and ANOVA (pRF number and visual field coverage) results comparing retinotopic properties across inversion and eccentricity. Bold text denotes statistical significance ($p < .05$).

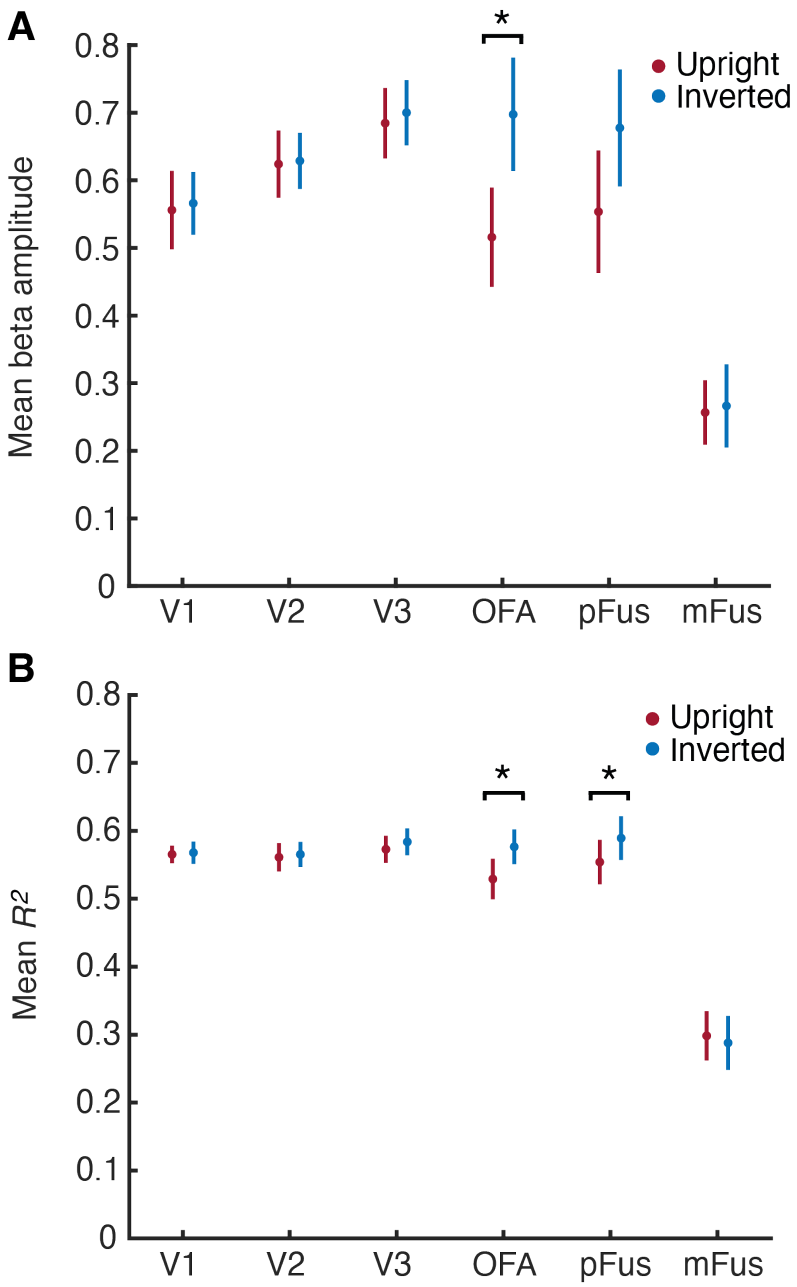


Figure S11. Mean beta amplitudes **(A)** and $R^2$ values **(B)** for upright (red) and inverted (blue) faces in all ROIs. Asterisks denote statistically significant differences ($p < .05$).
